# Altering the temporal regulation of one transcription factor drives sensory trade-offs

**DOI:** 10.1101/348375

**Authors:** Ariane Ramaekers, Simon Weinberger, Annelies Claeys, Martin Kapun, Jiekun Yan, Reinhard Wolf, Thomas Flatt, Erich Buchner, Bassem A. Hassan

## Abstract

Size trade-offs of visual versus olfactory organs is a pervasive feature of animal evolution. Comparing *Drosophila* species, we find that larger eyes correlate with smaller antennae, where olfactory organs reside, and narrower faces. We demonstrate that this tradeoff arises through differential subdivision of the head primordium into visual versus non-visual fields. Specification of the visual field requires a highly-conserved eye development gene called *eyeless* in flies and Pax6 in humans. We discover that changes in the temporal regulation of *eyeless* expression during development is a conserved mechanism for sensory trade-offs within and between *Drosophila* species. We identify a natural single nucleotide polymorphism in the cis-regulatory region of *eyeless* that is sufficient to alter its temporal regulation and eye size. Because Pax6 is a conserved regulator of sensory placode subdivision, we propose that alterations in the mutual repression between sensory territories is a conserved mechanism for sensory trade-offs in animals.

## INTRODUCTION

The senses animals rely on have been shaped during evolution to better navigate and exploit the environment. As a result, even closely related species living in different ecological niches show variation in the sizes and shapes of their sensory structures. Adaptive variation in visual sensory organs is a fascinating case in point. Darkness-adapted animals from phylogenetic groups as diverse as arthropods, fishes and mammals, often show loss of visual organs and an expansion of non-visual sensory structures, sometimes forming novel specialized organs, such as the ‘tactile eye’ of the star mole rat (Partha et al., 2017; Retaux and Casane, 2013). Conversely, arboreal, or tree-dwelling, rodents rely strongly on vision, and this behavioral trait is matched by the expansion of their primary visual cortical area (Campi et al., 2011; Campi and Krubitzer, 2010). In insects, the number of facets of the compound eye, known as ommatidia, can vary from a handful to several thousands. A large number of facets is associated with increased visual acuity and tends to be a distinctive feature of flying predator species such as dragonflies (Elzinga, 2003).

A striking, pervasive, yet poorly understood feature of natural variation in eye size is that it often occurs as a trade-off between visual and olfactory organs, where species with larger eyes have smaller noses and vice versa. A systematic analysis of sensory organ size across 119 mammalian species revealed a trade-off between nose size and eye size: arboreal species have relatively larger eyes and relatively smaller noses, while terrestrial mammals show the opposite trend (Nummela et al., 2013). Similarly, cave-dwelling fishes and arthropods have dramatic reduction in eye size and a reciprocal expansion in olfactory structures, such as olfactory pits or epithelia, or antennae in the case of arthropods (Retaux and Casane, 2013). These observations strongly suggest an obligate inverse relationship between the sizes of visual and olfactory organs. However, the genetic, molecular and developmental events that govern such trade-offs are essentially unknown.

Cellular and molecular mechanisms governing sensory development have been extensively studied in a variety of invertebrate (*D. melanogaster*, *C. elegans*) and vertebrate (mouse, chick, zebrafish) model organisms. These analyses have revealed that a common feature of sensory development is a shared developmental origin of most head sensory structures – such as eyes and noses – which derive from the subdivision of a single multipotent primordium. In vertebrates, most head sensory organs develop from a multipotent preplacodal ectoderm (Grocott et al., 2012; Singh and Groves, 2016). The olfactory and lens placodes derive from the subdivision of the anterior aspect of this common placode. Similarly, during *Drosophila* development, the ectodermal eye-antennal imaginal disc (EAD) gives rise to all external head structures, including the visual (compound eyes and ocelli) and olfactory (antennae and maxillary palps) sense organs, and the head cuticle. The subdivision of the EAD along the anterior-posterior axis forms the eye (posterior) and antennal (anterior) compartments, respectively. Each compartment is further subdivided into sensory organ versus head cuticle. Antagonistic relationships between gene regulatory networks (GRNs) and signaling pathways that promote different sensory identities regulate these processes (Grocott et al., 2012; Singh and Groves, 2016). For example, in vertebrates, the transcription factors (TFs) promoting olfactory and lens identity, Dlx5 and Pax6, are first co-expressed in the entire vertebrate anterior placode. Their expression domains segregate as the lens and olfactory territories become distinct. Ectopic expression of Dlx5 in the lens placode leads to the loss of Pax6 expression and of lens identity (Bhattacharyya et al., 2004). A similar mechanism subdivides the fruit fly EAD. At early developmental stages, TFs promoting eye fate, such as Pax6 homologs Eyeless (Ey) and Twin-of-Eyeless (Toy), and the SIX factor Sine oculis are ubiquitously co-expressed with TFs promoting antennal fate including the MEIS and PBX homologs Homothorax and Extradenticle. The eye and antennal TFs progressively segregate along the EAD’s anterior-posterior axis, delineating the posterior eye and anterior antennal compartments (Kenyon et al., 2003). Simultaneously, the antennal CUX-related TF Cut (Ct) appears in the anterior EAD, and progressively expands posteriorly as the expression of eye TFs like Ey retracts. Eye and antennal TFs mutually repress each other: Homothorax and Ct directly repress *ey* expression while Sine oculis represses *ct* (Anderson et al., 2012; Wang and Sun, 2012; Weasner and Kumar, 2013). Consequently, loss or gain of function of these selector TFs leads to the transformation of most of the head tissue into visual or olfactory organs at the expense of the other sensory structure (e.g. (Anderson et al., 2012; Czerny et al., 1999; Halder et al., 1995). In addition to specifying visual identity, Pax6 TFs also promote tissue growth both in vertebrates and in *Drosophila* (Zhu et al., 2017).

These observations show that the subdivision of a single multipotent primordium into distinct sensory territories through mutual repression by antagonistic TFs is a shared step of the development of head sensory organs across animals. It is thus tempting to speculate that evolutionary mechanisms have exploited this process leading to natural sensory size trade-offs between visual and olfactory organs. A hint in that direction comes from studies on *Astyanax* fishes, which live as cave or surface-dwelling morphs (Retaux and Casane, 2013). Cave morphs have small lenses and large olfactory placodes, while surface-dwellers show the reciprocal trade-off. Chemical manipulation of signaling pathways that regulate the subdivision of the lens versus olfactory territories mimics the differences observed between natural morphs (Hinaux et al., 2016). Whether this is a mechanism of natural variation in sensory trade-offs is unknown. Demonstrating a direct link requires the identification of naturally occurring causal genetic variants and the elucidation of their effect on the GRNs that regulate visual and olfactory sensory organ development. The paucity of model systems amenable to combining comparative, genetic, molecular and developmental analyses has thus far hindered such an endeavor.

We reasoned that natural variation in eye size between and within *Drosophila* species may offer precisely such a model. We therefore used comparative analyses combined with developmental, molecular and genome editing approaches to tackle this question. We show that differential subdivision of the EAD, resulting in different proportions of eye and antennal compartments, underlies eye size variation between and within *Drosophila* species. In both cases, this is associated with changes in the temporal regulation of the expression of *ey* during EAD subdivision. We also demonstrate that in *D. melanogaster* (*D. mel.)*, this is caused by a non-coding single polymorphic nucleotide (SNP) present in most natural populations of *D. mel*. This SNP is located in a putative repressor site for antennal factors within the eye enhancer of *ey*. Using CRISPR/Cas 9 genome editing, we show that this SNP is causal to facet number variation. Thus, changes in the subdivision of a multipotent sensory primordium, caused by subtle alterations of the mutual repression between distinct sensory fates, underlies natural variation in sensory trade-offs.

## RESULTS

### Reciprocal changes in the sizes of visual and non-visual head structures

The insect compound eye is composed of a crystalline array of small units, named facets or ommatidia. In insects, eye size depends both on the number and diameter of the ommatidia and is often negatively correlated with face and/or antenna size (Arif et al., 2013; Norry and Gomez, 2017; Posnien et al., 2012). In this study, we selected four *Drosophila* species, which presented a larger eye to head width ratio as compared to *D. mel*. (Figure 1A-1A’; Figure S1). We focused on *Drosophila pseudoobscura* (*D. pse.)*, which presented the largest difference in terms of ommatidia number, an increase of 35% as compared to *D. mel*. (Figures 1B and 1B’). In contrast, the two species presented no difference in facet diameter (Figures 1C and 1C’). We excluded that ommatidia number variation was a consequence of global variation in body size by measuring the scaling between facet number and two proxy measures for body size, thorax width and femur length (Figures S2A – S2B). Interestingly, in addition to a narrower face, the antennae of *D. pse*. were proportionally thinner as compared to *D. mel*. suggesting a trade-off between the size of compound eyes versus non-visual head structures (Figures 1D and 1E). Increased facet number has been associated with higher visual acuity in predator flies (Elzinga, 2003; Gonzalez-Bellido et al., 2011). We tested whether a modest variation such as the one observed between *D. pse*. and *D. mel*. was potentially relevant to visual function. We measured the minimal angular distance between two successive vertical black stripes resolved by the flies as a read-out of their visual acuity (Figure 1F) (Buchner, 1976; Gotz, 1964). *D. pse*. were able to distinguish between more closely juxtaposed stripes (minimal angle = 7.0°) as compared to *D. mel*. (minimal angle = 8.51°) (Figure 1F’). Thus, *D. pse*. have a better visual acuity correlating with an increased number of facets.

**Figure 1.**
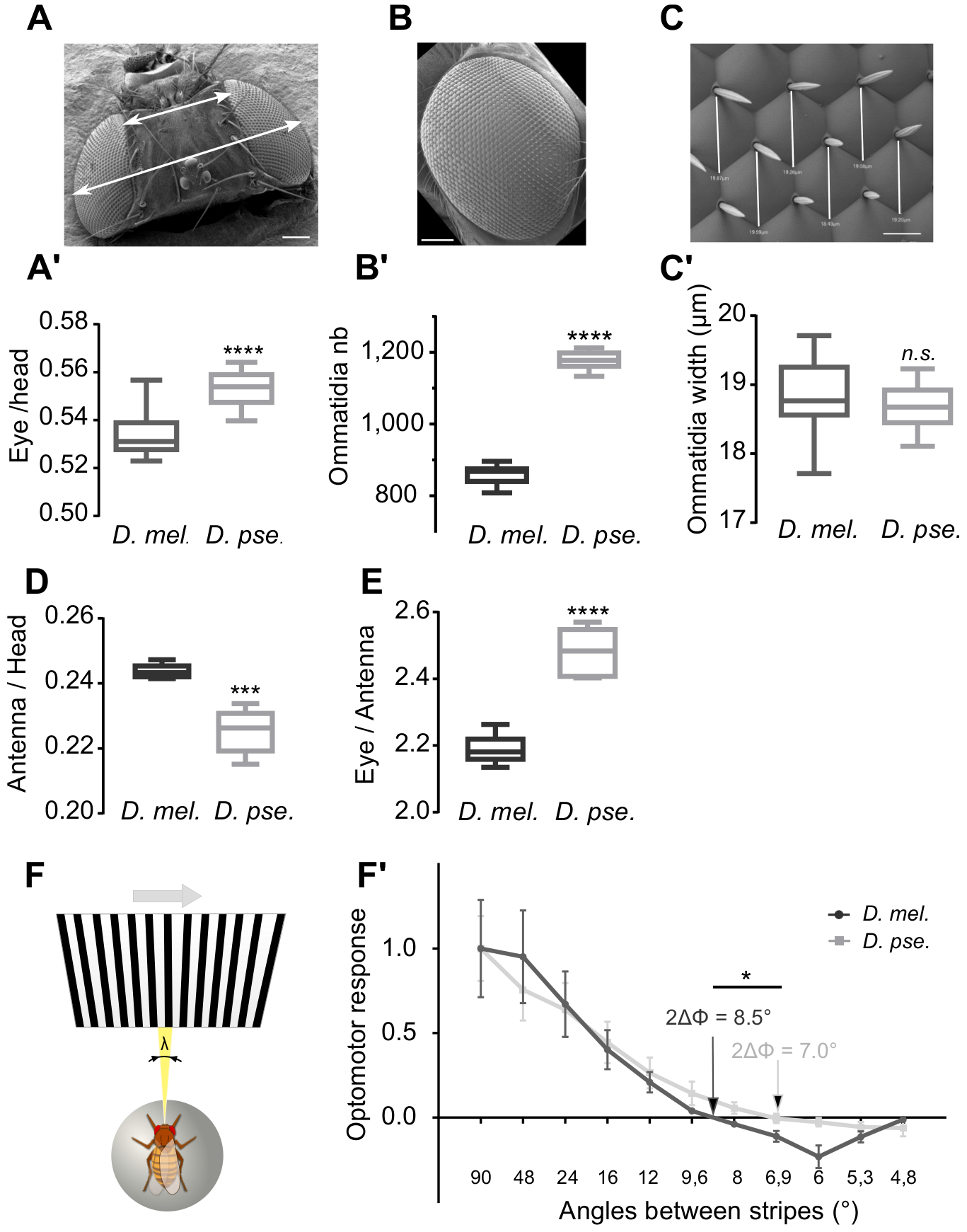
Natural variation in Drosophila facet number. (A to C’) Eye size comparisons between the two *D. mel*. (*D. mel*.) and *D. pse*. (*D.pse*.). Boxes indicate interquartile ranges, lines medians and whiskers data ranges (see also Figure S1). Scale bar for (A) and (B), 100 μm; for (C), 10 μm. (A and A’) Eye/head width ratio. Sample sizes: *D. mel*. (n=11), *D. pse*. (n=13). Two-tailed unpaired t-test: **** *p*<*0.0001*. (B and B’) Ommatidia number. Sample sizes: *D. mel*. (n=9), *D. pse*. (n=8). Two-tailed unpaired t-test: **** *p*<*0.0001*. (C and C’) Ommatidia width. Sample size: n=24. Two-tailed unpaired t-test with Welch’s correction: *n.s. p*=*0.1677*. (D) Antenna/head width ratio. Sample sizes: n=5. Two-tailed unpaired t-test: *** *p*=*0.0005*. (E) Eye/antenna width ratio. Sample sizes: n=5. Two-tailed unpaired t-test: **** *p*<*0.0001*. (E) Optomotor response set-up (see Materials and Methods). (E’) Optomotor response (normalized to max) of *D. mel*. and *D. pse*. females in function of stripe width (spatial wavelength λ); (mean ± sem). Arrows: spatial resolution (measured as the zero-crossing angle 2ΔΦ). Sample size: n=9. Two-tailed unpaired t-test: * *p* = *0.014*.

**TABLE 1:** Ommatidia numbers. Ommatidia numbers are counted on SEM images (count) or estimated from light-microscopy images using an ellipse-based method (estimated, see Materials & Methods and Figure S2). * Flies reared at 25°C.

|  | Ommatidia nb (*count*) |  | Ommatidia nb (*estimated*) |  |
| --- | --- | --- | --- | --- |
|  | Mean | St Dev | Mean | St Dev |
| Canton-S | 869.8* | 17.96* | 828.0 | 34.00 |
| Hikone-AS | 742.4* | 20.17* | 736.0 | 27.09 |
| *D. pse.* | 1177* | 28.02* |  |  |
| F1 CS/Hik |  |  | 774.6 | 34.77 |
| *ct*<sup>5687</sup>/*ct*-GAL4 |  |  | 835.7 | 25.88 |
| *ct*<sup>5687</sup>/TM3,Sb |  |  | 802.9 | 26.99 |
| *ct*<sup>4138</sup>/*ct*-GAL4 |  |  | 818.6 | 23.24 |
| *ct*<sup>5687</sup>/TM3, Sb |  |  | 777.6 | 22.99 |
| Canton S BH |  |  | 830.8 | 36.42 |
| Canton S TP |  |  | 775.4 | 21.69 |
| Canton S RD |  |  | 750.4 | 20.26 |
| CRISPR control | 713.8 | 29.77 | 745.1 | 43.76 |
| CRISPR 1 | 737.0 | 21.12 | 791 | 44.49 |
| CRISPR 2 |  |  | 785.1 | 35.89 |
| CRISPR 3 |  |  | 758.2 | 43.67 |
| CRISPR 4 |  |  | 770.8 | 31.63 |

### A trade-off between eye and non-eye progenitor fields

What is the developmental origin of facet number variation between *D. mel*. and *D. pse*. and why does it inversely correlate with the size of non-visual structures? All external structures of the head of the adult fly including the sensory organs develop from the EAD (Figure 2A). The eye field occupies most of the posterior EAD compartment and is marked by the expression of Eyes Absent (Eya; Figure 2B) (Roignant and Treisman, 2009). We measured the surface of the eye field in late, fully-grown, EADs (stage P0) using Eya and found that the eye field was 31% larger in *D. pse*. than in *D. mel*. (Figure 2B’). This is very close to the 35% difference in the number of adult eye facets. We therefore queried the developmental origin of the difference in eye field size between the two species, and considered several possibilities. A first possibility is that the initial pool of embryonic cells forming the EAD differs between the two species. In the late embryo (stage 17), the EADs is composed of a few dozen closely juxtaposed cells located anterior to the brain. Using Ey as a marker, we quantified and compared the number of embryonic EAD cells between the two species (Figures 2D and 2D’) but found no significant difference in the number of EAD progenitors, ruling out this first possibility. Variation in eye field size could also originate from different rates of proliferation. However, the similar density of mitotic cells in the proliferating eye field in EADs of the two species did not support this hypothesis (Figures 2F and 2F’). In addition, the density of ommatidia progenitor cells in the eye field, characterized by the expression of the proneural factor Atonal, was similar between the two species (Figures S3A-S3A”). Finally, variation in eye field size could also derive from a change in the subdivision of the EAD between eye and non-eye fields. To test this possibility, we compared the proportion of the EAD occupied by the eye field in early L3 imaginal discs, after the subdivision between the fields is completed (Figure 2E). The total EAD size was similar between the two species, confirming that it underwent similar growth during prior larval development (Figure 2E’). In contrast, already at this early stage, the eye field was proportionally larger in *D. pse*. than in *D. mel*. (Figure 2E”). Thus, the two species differ by the proportion of the multipotent EAD dedicated to the eye versus non-eye tissues, resulting in different proportions of the head structures in the adult. Therefore, the species variation in eye size involves a developmental trade-off between eye and non-eye primordia.

**Figure 2.**
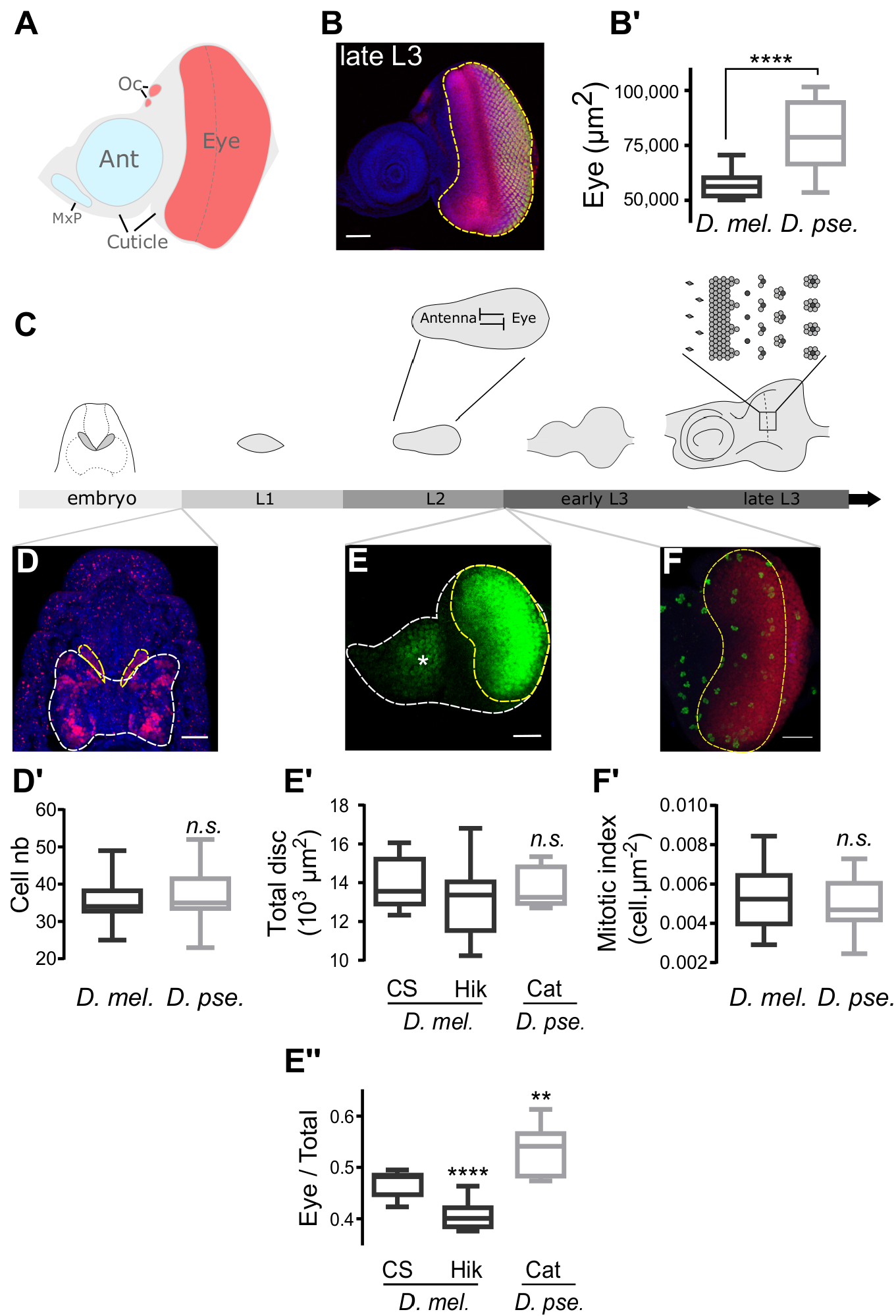
Developmental origin of eye size variation in *D. mel*. and *D. pse*.. (A) Schematics of the distinct fields of the EAD giving rise to the adult sensory and non-sensory head structures (Ant, antenna; MxP, maxillary palps; Oc, ocelli). Anterior is at the left. (B) Late L3 EAD from *D. pse*.. The eye field (yellow dashed line) is labelled with anti-eya (red) and committed photoreceptors are shown in yellow (anti-elav); blue: DAPI. Anterior is at the left. Scale bar: 50 μm. (B’) Eye field surface in late (P0) eye-antennal discs. Sample sizes: *D. mel*. (n=19); *D. pse*. (n=19). Two-tailed unpaired t-test with Welch’s correction **** *p*<*0.0001*. (C) Schematics of eye-antennal disc development. Insets: (C’) segregation of “antennal” and “eye” compartments during L2; (C”) determination of ommatidia precursor cells during L3. (D) Dorsal view of a late Canton-S embryo depicting the Ey-positive EADs (yellow dashed line) flanking the brain (white dashed line). Anterior is at the top. Scale bar: 25 μm. (D’) Cell content of late embryonic EADs. Sample sizes: *D. mel*. (n=22); *D.pse*. (n=33). Two-tailed unpaired t test; *n. s. p*=*0.0959*. (E) Early L3 Canton-S EAD co-labelled by Eya (yellow dashed line) and Ct (asterisk). Anterior is at the left. Scale bar: 25 μm. (E’) Total surface of the EADs (in μm^2^). Ordinary one-way ANOVA (*p*=*0.4520*) followed by Dunnett’s multiple comparisons tests. (E”) Ratio of eye field *vs* total EAD surface. Ordinary one-way ANOVA (*p*<*0.0001*) followed by Dunnett’s multiple comparisons tests. Sample sizes: CS (n=11); Hik (n=9); Cat (n=8). (F). Proliferative portion of the eye field (dashed yellow line). Red: eye field (anti-Eya); Green: mitotic cells (anti-phosphorylated Histone 3). Anterior is at the left. Scale bar: 25 μm. (F’) Mitotic index of the proliferative eye field in *D. mel*. and *D. pse*. early L3 EADs. Sample sizes: *D. mel*. (n=20); *D. pse*. (n=8). Two-tailed unpaired t-test *n.s. p*=*0.6185*. Boxes indicate interquartile ranges, lines medians and whiskers data ranges in all charts (B’, D’, E’, F’).

### Temporal regulation of EAD subdivision governs the trade-off between eye and non-eye fields

**Figure 3.**
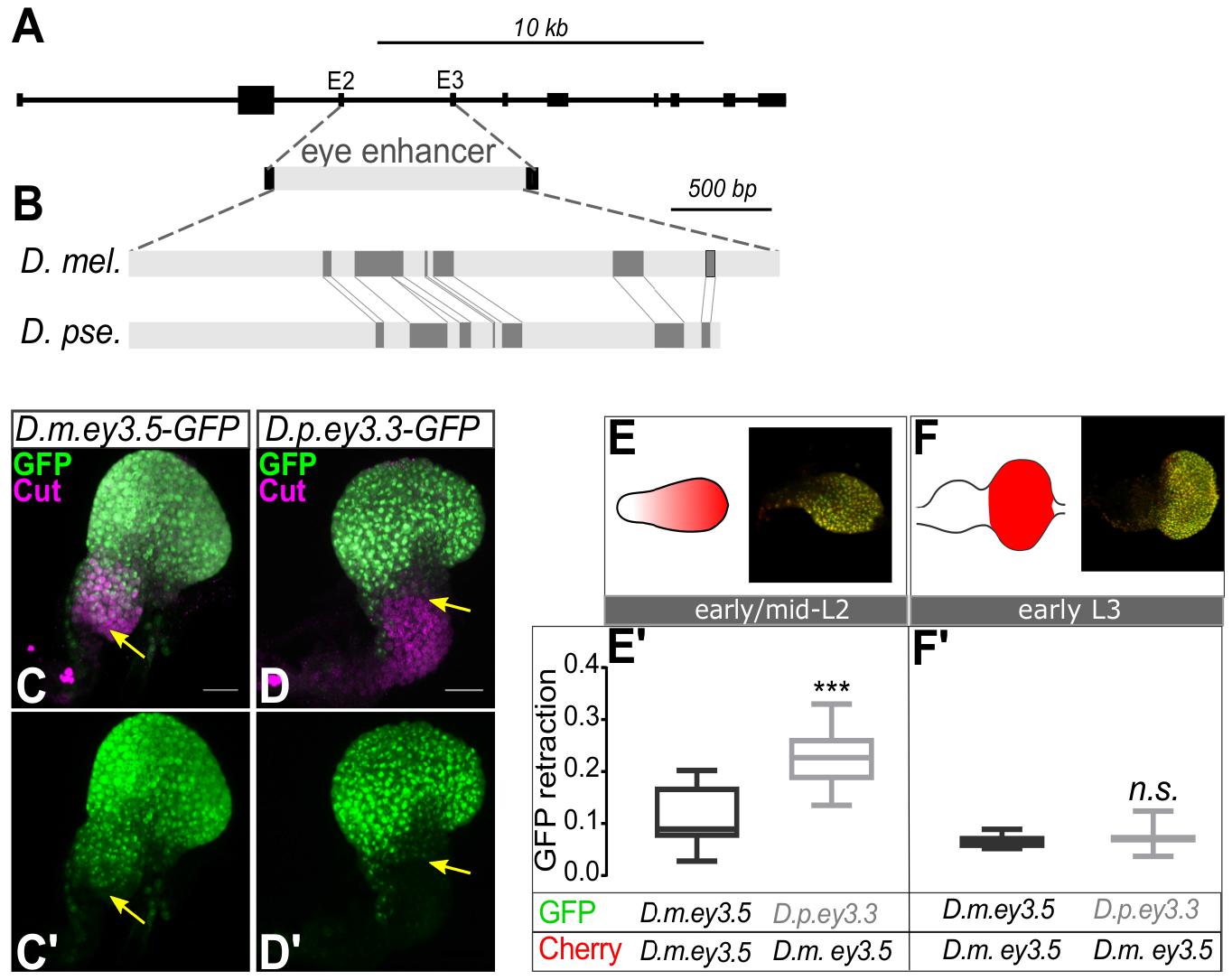
Different temporal regulation of EAD subdivision governs the trade-off between eye and non-eye fields. (A) Structure of the *ey* locus showing the location of the intronic *ey*^*3.5*^ eye enhancer (Hauck et al., 1999). E2: exon 2; E3: exon 3. (B) Alignment of the *ey* eye enhancer between *D. mel*. and *D. pse*.. Dark grey boxes represent the fragments of the *D.mel*. intron sequence that aligned with the orthologous region of *D. pse*. using BLAST. (C to D’) GFP expression driven by *D. mel. ey3.5* and *D. pse.ey3.3* eye enhancers in mid-L2 EADs counterstained with anti-Cut. *Green*: GFP; *magenta*: anti-Cut. Yellow arrows indicate the anterior limit of GFP expression. Scale bars: 20 μm. Anterior is at the bottom. (E and F) Schematics and immunostainings showing ongoing (E, early / mid-L2 stage) and full (F, early L3 stage) posterior retraction of *ey* enhancer activity during EAD development. (E’ and F’) Pairwise comparison of the difference in expression between GFP and mCherry, measured as the proportion of mCherry that was not colocalized with GFP, when driven by distinct combinations of *ey* eye enhancers during (E’, early/mid-L2 stage) and after (F’, early L3 stage) posterior retraction of ey enhancer activity. (*D.m.ey3.5, D. mel. ey* enhancer; *D.p.ey3.3*, *D. pse*. enhancer). (E’) Early/Mid-L2 stage. Sample sizes (n=8, n=12). Two-tailed unpaired t-test *** *p*=*0.0001*. (D’) Early-L3 stage. Sample sizes (n=10, n=3). Mann-Whitney test *n.s. p= 0.6643*

What are the regulatory mechanisms governing this sensory organ size trade-off? EAD subdivision requires the temporally progressive restriction of selector TFs expression to the anterior ‘antennal’ or posterior ‘eye’ compartments, a process completed by mid to late second instar larval stage (L2) (Kenyon et al., 2003). At this developmental time point, the mutually exclusive expression domains of antenna and eye selectors define the relative sizes of the compartments. In *D. mel*., a 3.2 kb *cis*-regulatory intron governs *ey* expression during eye development (Figure 3A). We cloned the orthologous intron from *D. pse*. based on the conservation of the flanking exons. The *D. pse*. intronic sequence is slightly shorter (3.0 kb) with 22% of the intron from *D. mel*. aligning to the corresponding sequences in *D. pse*. (Figure 3B). Nonetheless, when inserted at the same position in *D. mel*. genome, both *D. mel*. and *D. pse*. introns were able to drive GFP expression in the EAD throughout eye development, revealing global functional conservation (not shown).

We tested whether, despite their overall functional conservation, subtle changes in *ey* regulation exist between *D. mel*. and *D. pse*. introns (Figures 3C-3D’). At mid-L2 stage, we noted that GFP expression driven by the *melanogaster* enhancer (*D.m.ey3.5*) (Figures 3C and 3C’) extended further anteriorly into the antennal compartment as compared to the *pseudoobscura* enhancer (*D.p. ey3.3*) (Figures 3D and 3D’). This means that the posterior retraction of expression driven by the two *ey* enhancers occurs at different velocities. To quantify this effect, we generated two lines of transgenic *D. mel*. flies. The first line carries two transgenes driving the expression of the red fluorescent protein mCherry and of the green fluorescent protein (GFP), respectively, both under the control of *D. mel. ey* enhancer. In this control line, any difference in the expression of mCherry and GFP driven by the same enhancer must only be caused by different dynamics of the two fluorescent proteins. In the second line, mCherry was driven by the *D. mel. ey* enhancer, while GFP was driven by *D. pse. ey* enhancer. In this case, the differences in expression between the GFP and mCherry is caused both by different dynamics of the fluorescent proteins as well as differences in their transcriptional regulation. Thus, to detect differences in the activity of *D. mel*. and *D. pse. ey* enhancers, we performed pairwise comparisons of the difference between GFP and mCherry expression in line 1 versus line 2. In early L3 discs, when the antennal and eye compartments have already segregated, the co-expression of mCherry and GFP driven by either *D. mel*. or *D. pse*. regulatory sequences were indistinguishable (Figures 3F and 3F’). In contrast, at mid L2, during the process of *ey* retraction, the posterior retraction of the GFP was more posteriorly advanced when driven by the *D. pse*. than by the *D. mel*. enhancer (Figures 3E and 3E’). Therefore, the partitioning of the EAD into eye and non-eye fields occurs at an earlier time point in *D. pse*. compared to *D. mel*.. Since Ey positive cells proliferate more than Ey negative cells (Zhu et al., 2017), earlier establishment of the two sensory fields would drive greater differential growth.

**Figure 4.**
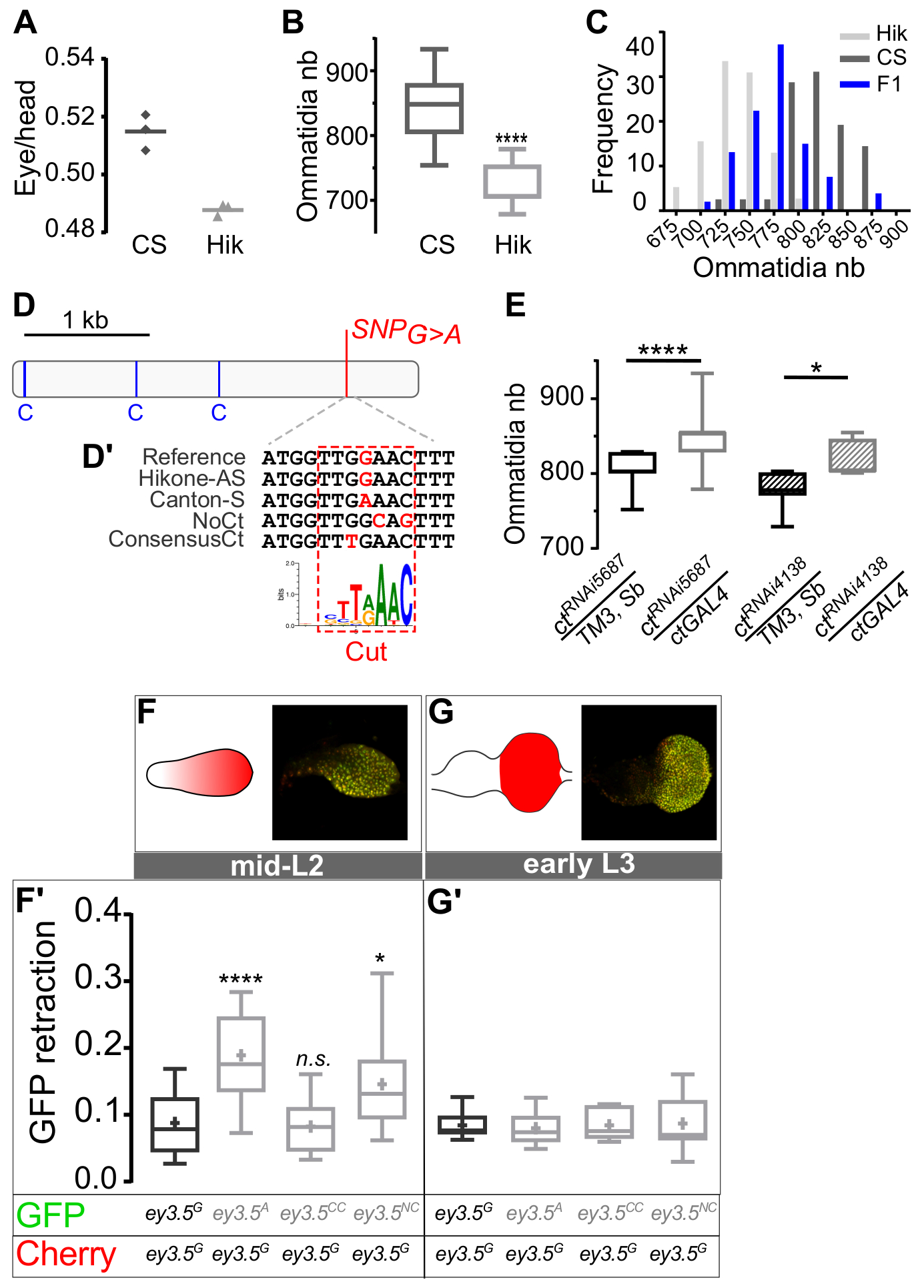
Developmental and regulatory origin of intraspecific eye size variation. (A) Eye/head width ratio in Canton-S (CS) and Hikone-AS (Hik). Also see Figure S1. (B) Ommatidia number. Sample sizes: Canton-S (n=15), Hikone-AS (n=16). Two-tailed unpaired t-test: **** *p*<*0.0001*. (C) Frequency distribution of ommatidia numbers of Canton-S (n=42), Hikone-AS (n=39) and their F1 progeny (n=54). (D) Schematics of the *D. mel. ey* eye enhancer (*ey*^*3.5*^) showing the localization of the *G*>*A* substitution (chr4: 710326) and of three binding-sites for Ct (C, blue lines) (Wang and Sun, 2012). (D’) Sequence flanking the *G*>*A* substitution in natural and synthetic enhancer alleles. The dashed red rectangle delineates the putative Ct binding site (Zhu et al., 2011). (E) Ommatidia number variation following RNAi-mediated knock-down of *ct*. Sample sizes from left to right (n=13; n=23; n=10; n=8). Ordinary one-way ANOVA **** *p*<*0.0001* followed by Sidak’s multiple comparisons: **** *p*<*0.0001*, \**p*=*0.0126*. See also Figure S4. (F to G’) Pairwise comparison of the expression of the four *D. mel. ey* enhancer alleles: *ey3.5*^*G*^ (Hikone-AS); *ey3,5*^*A*^ (Canton-S); *ey3.5*^*CC*^ (ConsensusCt); *ey3.5*^*NC*^ (NoCt). (G and H) Schematics and immunostainings of EADs with ongoing (F, early/mid-L2 stage) and full (G, early L3 stage) posterior retraction of ey enhancer activity.

### A conserved mechanism of sensory trade-offs

To understand the genetic basis of sensory trade-off in *Drosophila*, we exploited the fact such trade-offs have also been observed within single fly species (Arif et al., 2013; Cowley and Atchley, 1990; Norry and Gomez, 2017; Posnien et al., 2012). We found that two wild-type *D. mel*. laboratory strains, Canton-S and Hikone-AS, show differential eye size and eye-to-face ratio associated with changes in ommatidia number (12, 5 % more facets in Canton-S) and diameter (Figures 4A and 4B, Figures S1A - S1D, Table 1). F1 progeny of a cross between Canton-S and Hikone-AS presented intermediate ommatidia numbers relative to their parents, demonstrating the heritable nature of the trait (Figure 4C). We asked if facet number variation between Canton-S and Hikone-AS also originate from changes in the subdivision of the EAD into eye versus non-eye territories. We compared the subdivision of early L3 EADs between the two strains. While the size of the entire EAD was unchanged, the eye field was proportionally larger in Canton-S than in Hikone-AS (Figures 2E – 2E”). These data suggest that, despite 17 - 30 million years of separated evolution between the two species groups (Obbard et al., 2012), ommatidia number variation between *D. mel*. and *D. pse*. and between two *D. mel*. strains, shares a common developmental logic.

### A single nucleotide in a Ct binding site distinguishes the Canton-S and Hikone-AS *ey* regulatory sequences

Does the difference in EAD subdivision between the Canton-S and Hikone-AS also result from a differential temporal regulation of *ey*? To answer this question, we cloned and aligned the *ey cis*-regulatory sequence from the Canton-S and Hikone-AS strains (Figure 4D). In contrast to the significant divergence observed between *D. mel*. and *D. pse*., Hikone-AS and Canton-S intron sequences were nearly identical and differed only by a single nucleotide over the entire 3.2 kb intronic region. The polymorphism is a Guanine (*G*) in Hikone-AS to Adenine (*A*) in Canton-S substitution and is located in a sequence resembling a Ct binding site, which is distinct from three Ct binding sites previously described in the *ey cis*-regulatory sequence (Figure 4D) (Wang and Sun, 2012). This suggests that the degree of *ey* repression by Ct, a selector TF for antennal fate, may mediate facet number variation (Wang and Sun, 2012; Weasner and Kumar, 2013). Consistent with this, RNAi knock-down of *ct* expression during EAD development was sufficient to increase facet number in the adult eye (Figure 4E, Figure S4).

These findings raise two questions: first, is the *G* to *A* substitution in the *ey cis*-regulatory sequence sufficient to cause temporal changes in its activity; and second, if so, might such changes be caused by alterations in the regulation of the *ey* enhancer by Ct? To tackle these two questions, we used the same strategy described above for comparing the *D. pse*. and *D. mel*. enhancers using GFP and mCherry reporters. We first compared the activities of the Canton-S (*A*-allele) and the Hikone-AS (*G*-allele) of the *ey 3.5 cis*-regulatory sequences. At early/mid L2, during EAD subdivision, mCherry and GFP co-expression differed between the alleles such that the *A*-carrying variant (Canton-S; larger eyes) showed further posterior retraction of GFP expression than the *G*-carrying variant (Hikone-AS; smaller eyes) (Figure 4F-4F’). In contrast, at early L3, after EAD subdivision is completed, the two alleles drove similar expression of GFP and mCherry (Figure 4G-4G’).

Could this differential temporal retraction of the *ey* enhancer be caused by changes in ey repression by Ct, as suggested by the Ct RNAi results? To test this, we created two new synthetic *ey* enhancers (Figure 4D). The first, which we call the *NoCt* variant, is predicted to abolish Ct binding to the site harboring the *G/A* SNP. The second, which we call *ConsensusCt*, creates a Ct consensus-binding motif at that position. Remarkably, the *ConsensusCt* variant behaved similarly to the *G*-allele, while the *NoCt* variant mimicked the *A*-allele in that it caused faster posterior retraction of *ey* enhancer activity (Figure 4F-G’) suggesting that the Canton-S *A* allele may constitute a lower affinity site for the Ct repressor as compared to the Hikone-AS *G* allele.

Thus far, we showed that the changes in EAD subdivision between Hikone-AS and Canton-S and between *D. mel*. and *D. pse*. are both driven by differential temporal dynamics of the posterior retraction of *ey* expression. Within the *D. mel*. this is associated with a single substitution in *ey* eye enhancer, which introduces subtle changes in *ey* regulation, likely by affecting its repression.

### A common SNP in *D. mel*. natural populations is associated with facet number variation

Because Hikone-AS and Canton-S flies have been in artificial lab culture conditions for decades, we asked if either of these two alleles is actually found in natural fruit fly populations. By investigating allele frequency patterns in whole-genome data of worldwide population samples, we found that most natural populations from Europe, North America, Asia and Australia are polymorphic at this position. Thus, neither of the two alleles corresponds to a *de novo* mutation and variation at this position corresponds to a relatively frequent SNP (Figures 5A and A’; Table S2). Populations from Sub-Saharan Africa are mostly fixed for the *G*-allele suggesting that the *A*-allele is a derived variant that appeared after *D. mel* left Africa and colonized the rest of world. In line with this hypothesis, we found statistical evidence (FET test, p=0.02) that the few African populations carrying the *A*-variant are more likely to be admixed with European genetic variation than the ones with the putatively ancestral *G*-allele (Table S3). Moreover, the frequency of the *A*-variant decreased from West to East in European populations (Figure 5A’ and 5B). The slope of the longitudinal frequency cline of the *ey* SNP deviated significantly from that of 21,008 genome-wide SNPs in short introns that presumable evolved neutrally (Figure 5C), suggesting that the clinal pattern is not solely the result of neutral evolution or demography (see also Figure S5).

**Figure 5.**
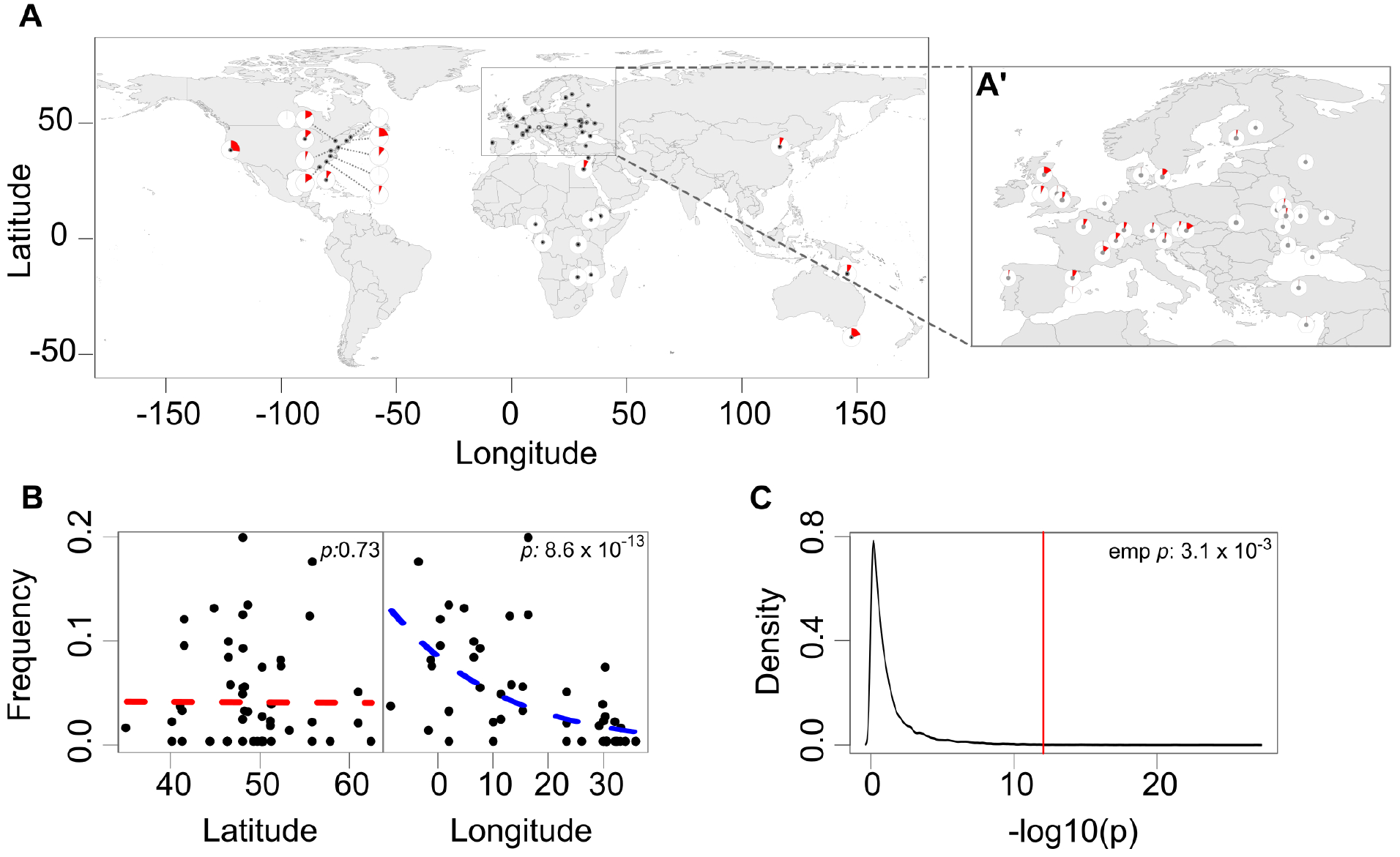
Worldwide distribution of the *G*>*A* substitution (chr4: 710326) in the genome of *D. mel*. natural populations. See also Figure S5. (A) Worldwide allele frequency patterns. Frequencies of the *A*-variant are shown as red areas of the pie charts for worldwide populations with a sample-size ≥ 10 (see tables S2 and S3 for details). The exact geographic location for each samples is indicated by a black dot. (A’) Frequency distribution of the *A*-variant in Europe revealing the longitudinal clinal distribution of the allele. (B) Scatter plots showing allele frequencies of the A-variant along the latitudinal (left box; red regression curve) and the longitudinal axis (right box; blue regression curve) in Europe. The top-right *p* values show the significance from generalized linear models (GLM). (C) The line plot shows the empirical cumulative density function (ECDF) from −log_10_ transformed *p* values of GLMs with longitude as the predictor variable for 21,008 neutrally evolving intronic SNPs (black curve). The vertical red line highlights the −log_10_ *p* value for the focal SNP (*p*=*8.6*× *10*^−*13*^) which is inferior to the ones of 99.69 % of the neutral SNPs, indicating that it stronger correlation with longitude. The empirical *p* value (*p*=*0.0031*) is calculated from the area confined by the *p* value of 4:710326 and the tail of the ECDF.

### Causal effect of the SNP

We further noted that natural populations from North-East America, where the Canton-S strain originated, are highly polymorphic for the *ey* SNP (Figure 5A, Table S2). By comparing Canton-S flies from three laboratories, we discovered that, while our Canton-S lab isolate (henceforth Canton-S^BH^) carries the *A*-allele, two other strains from two different laboratories in Paris, France (T. Préat) and Florida, USA (R. Davis) were homozygous for the *G*-allele, similar to Hikone-AS. This strongly suggests that the original Canton-S population was polymorphic and that the two alleles were eventually segregated during the separate maintenance of different laboratory stocks (Colomb and Brembs, 2014). This provided a unique opportunity to quantify the contribution of the *G/A* SNP to eye size in a relatively homogenous background. By comparing ommatidia numbers between the three stocks, we observed that Canton S^BH^ flies have significantly more facets than its two siblings (Figure 6A and Table 1). These data suggest that the *A*-variant may be sufficient to drive larger facet numbers. To test this idea, we used CRISPR/Cas9 to introduce the *A*-allele in a *G*-homozygous stock. We recovered one transformant male carrying the *A*-allele and no other mutations in the *ey* regulatory intron and established four separated stocks from its progeny to account for potential subtle genetic background effects. The original *G*-homozygous stock underwent the same crossing scheme and was subsequently used as control. Comparing the five lines revealed an increased mean facet number for all four engineered *A*-carrying stocks as compared to the *G*-homozygous controls, reaching statistical significance in three out of the four *A*-homozygous strains (Figure 6B, Table 1). The *G*>*A* substitution recapitulates up to 49 % of the difference between Hikone-AS and Canton-S^BH^ and up to 86% of the variation observed between the genetically Canton-S^BH^ and Canton-S^RD^ and Canton-S^TP,^ respectively. Thus, the *ey cis*-regulatory SNP is causal to facet number variation.

**Figure 6.**
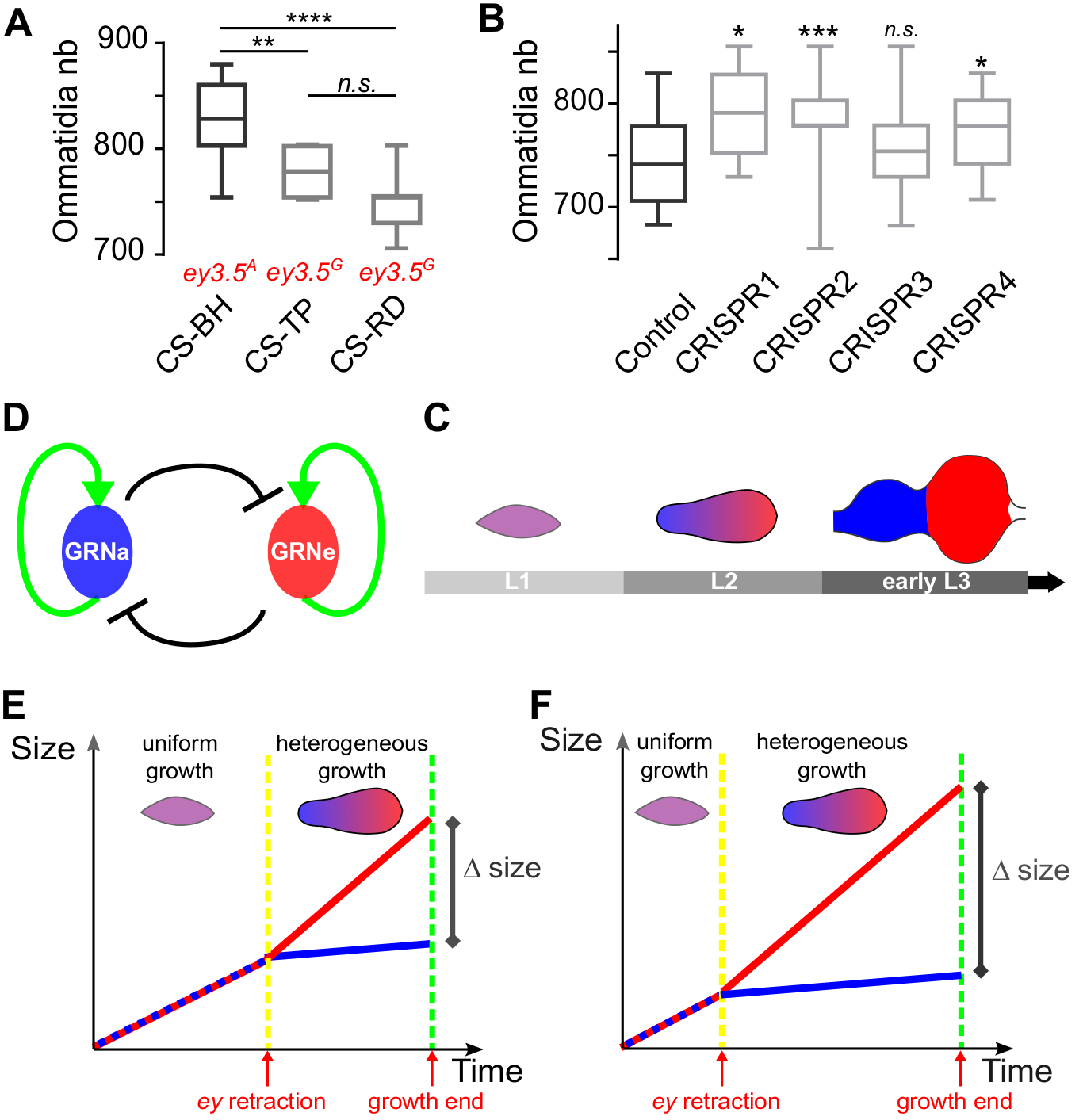
The non-coding *G*>*A* SNP in *ey* enhancer causes facet number variation. (A) Ommatidia numbers in three Canton-S strains with different *ey* SNP alleles (in red). Sample sizes: Canton-S^BH^ (CS-BH, n=18), Canton-S^TP^ (CS-TP, n=16), Canton-S^RD^ (CS-RD, n=19); Kruskal Wallis test **** *p*<*0.0001* followed by Dunn’s multiple comparisons: ** *adjusted p*=*0.0041*; **** *adjusted p*<*0.0001*; *n.s. adjusted p*=*0.1180*. (B) Ommatidia numbers in CRISPR *A*-variant and control *G*-variant homozugous fly eyes. Sample sizes: from left to right (n=24, n=8, n=32, n=33, n=45); Ordinary one-way ANOVA *** *p*=*0.0009* followed by Dunnet’s multiple comparisons between the control and the four CRISPR lines. (A and B) Boxes indicate interquartile ranges, lines medians and whiskers data ranges. (C to F) Model of the developmental origin of the trade-off between EAD derived structures. (C) The subdivision of the EAD into an anterior (“antennal”) and posterior (“eye”) compartments involves a bistable switch by which GRNs promoting eye (GRNe, *in red*) and antennal (GRNa, *in blue*) identity act antagonistically by activating their own and repressing the alternative GRN’s activity. (D) The bistable switch between the GRNa and GRNe results in the progressive segregation of the expression domains of TFs promoting eye (posterior, *in red*) versus antennal (anterior, *in blue*) fates during successive stages of EAD development. Anterior is on the left. (E and F) Our model, inspired from Slatkin (Slatkin, 1987), proposes that different temporal dynamics of the posterior retraction of *ey*, a promotor of EAD proliferation, by changing the relative duration of uniform versus heterogeneous growth, modifies the proportion between the antennal and eye compartments. This could be caused by genetic changes affecting the dynamics of the bistable switch between GRNe and GRNa, like in the case of the ey G>A substitution.

## DISCUSSION

In 1987, Montgomery Slatkin proposed a mathematical model (Slatkin 1987), which he referred to as “unrealistically simple”, that addresses “the case of a bifurcating sequence of events in which traits develop from the same tissue until a transition occurs, after which they develop partially independently”. This model predicts that mutations modifying the time at “which this transition takes place will change the relative size of the traits. Sensory trade-offs between visual and olfactory sensory organs are pervasive in nature and represent precisely the types of traits referred to in Slatkin’s model. However, whether and how visual-olfactory sensory trade-offs follow a “Slatkin model” and if so, what the mechanistic biological basis of such a model remained unexplored.

In this study, we identify a mechanism explaining the pervasive trade-off between visual, olfactory and non-sensory head structures using the *Drosophila* genus as a model. We find that differential subdivision of the head primordium into distinct progenitor fields during development is at the origin of this trade-off. We further demonstrate that this is associated with differential temporal regulation of the expression of the eye field selector transcription factor Ey. In the fruit fly, Ey promotes cell proliferation in the head primordium (Zhu et al., 2017). We propose a model (Figures 6D - 6F) whereby early in development, the homogenous expression of *ey* causes homogenous growth throughout the entire EAD. Later, the progressive retraction of *ey* expression from the anterior antennal compartment creates an asymmetry in growth rate. Modulating the velocity of *ey* retraction through mutations affecting the bistable switch between GRNs governing antennal *vs* eye identity, changes the relative time during which the anterior and posterior compartments grow different rates resulting in their different proportions. This provides biological evidence for mathematical models linking heterochrony in development to changes in adult traits (Cowley and Atchley, 1990; Riska, 1986; Slatkin, 1987).

The same developmental and regulatory features – temporal changes in the subdivision of the head primordium and in *ey* regulation – underlie eye size variation between *D. mel*. and *D. pse*., and within *D. mel*.. Due to the tight coordination between growth, specification and patterning processes, and because the eye shares a common primordium with other head sensory and non-sensory structures, most developmental changes likely result in dramatic “pleiotropic” effects, leaving little room for non-pathological variation. In this context, changing the time of onset of the subdivision of the head primordium into distinct territories may constitute a safer, and hence preferred, route to size variation. This mechanism may be a conserved determinant of sense organ size variation, thereby explaining the pervasiveness of trade-offs between olfactory and visual organs.

Within *D. mel*., a naturally occurring SNP in the eye-enhancer of *ey* is sufficient to modulate the velocity of the posterior retraction of the enhancer activity and to vary facet numbers in the adult eye. The SNP, a *G/A* substitution, is located in a putative binding site for *ey* repressor Ct suggesting that changes in *ey* repression by Ct are responsible for changing the dynamics of *ey* regulation and ultimately for causing morphological variation. This view is further supported by our findings that (i) synthetic mutations predicted to support or reduce Ct binding mimic the effect of the SNP on the velocity of the enhancer retraction; and that (ii) knocking-down Ct expression increase compound eye size. Turning *G*-carrying into *A*-carrying flies using CRISPR/Cas9 recapitulated up to 49% of the natural variation observed between Canton-S and Hikone-AS but up to 86% of the variation between two genetically highly similar Canton-S lab strains (Figure 6A, Table 1). This indicates that additional genetic changes must contribute to eye size variation between Canton-S and Hikone-AS. Analyses of the genetic origin of natural variation in embryonic trichome pattern or abdominal pigmentation in fruit flies identified several synergistic mutations in a single enhancer, each alone contributing a small effect to the trait (Frankel et al., 2011; McGregor et al., 2007; Rebeiz et al., 2009). In contrast, the Canton-S and Hikone-AS *ey* eye enhancers differ by only this one SNP, indicating that other factors lie outside from this region. We predict that these other mutations may target loci governing, either in *cis* or in *trans*, the velocity of the segregation between the antennal and eye GRNs.

In vertebrates, the regulation of the development of the lens and olfactory placodes shares significant commonalities with *Drosophila* sense organ development (Grocott et al., 2012; Singh and Groves, 2016). Their common primordium, the anterior placodal region, expresses Pax6. In the absence of Pax6, both the lens and olfactory placodes fail to thicken and to develop properly (Ashery-Padan et al., 2000; Collinson et al., 2000; Quinn et al., 1996). The onset of the activation of a lens placode Pax6 enhancer can be regulated by manipulating the affinity of two suboptimal binding sites for its activator pKnox1(or Prep1), a Meis related factor (Rowan et al., 2010), providing a plausible hypothesis for how lens size might vary between vertebrates. In the fish, *Astyanax mexicanus*, cave and surface morphs differ in the proportion between lens and olfactory placode size and manipulating signaling pathways involved in anterior placode subdivision phenocopies the differences between the natural variants (Hinaux et al., 2016). Thus, similar to *Drosophila*, mutations affecting the subdivision of the anterior placodal region into lens and olfactory territories could generate antagonistic changes in lens and olfactory epithelium size. We therefore hypothesize that natural occurring genetic variants acting either in cis or trans on Pax6 regulation is a universal mechanism for the regulation of the trade-offs between visual and olfactory organs.

The global distribution of the *G/A* SNP in *D. mel*. natural populations, based on the analysis of currently available genomes, suggests that the *A*-allele is a derived variant that appeared after *D. mel*. migrated out of Africa. In Europe, the SNP shows a longitudinal gradient of variation, which cannot be solely explained by demographic effects, suggesting a potential adaptive value to the sensory trade-off. The difference in visual acuity between *D. mel*. and *D. pse*. raises the obvious question of the nature of such a putative adaptation. However, facet number is one of the traits affected by the trade-off in head primordium subdivision. Therefore, the target of selection could equally be the size of the antenna, the distance between the two eyes or any combination thereof.

In “The Origin of Species”, Charles Darwin referred to the evolution of the eye as a challenge to his theory (Darwin, 1872). He also discussed the importance of correlation between body parts concluding that it was “most imperfectly understood”. During the last decades, the common origin of animal eyes and their evolution over long evolutionary distances has been abundantly documented (Gehring, 2014). However, the developmental mechanisms by which small-scale variation in eye size or shape can take place without disrupting its organization and function remain largely elusive (Dyer et al., 2009). We have demonstrated that a single nucleotide change in a core regulator of eye development is sufficient to generate reciprocal sensory organ size variation, potentially providing a quick route to behavioural changes, and perhaps adaptation. As predicted by Darwin, adaptive variation in head derived structures, including the eye, can be produced by the accumulation of modest morphological changes, which our data suggest may be caused by a small number of genetic variants affecting the temporal regulation of core developmental networks.

## METHODS

### Contact for reagent and resource sharing

Further information and requests for resources and reagents should be directed to and will be fulfilled by the Lead Contact, Bassem Hassan.

### Fly stocks and husbandry

Fly stocks used in this study are: *D. melanogaster*: Canton-S^BH^ (Hassan lab); Canton-S^TP^ (Préat lab, provided by P. Callaerts); Canton-S^RD^ (Davis lab, provided by P. Callaerts); Hikone-AS (Kyoto DGRC 103421); DGRP-208 (*D.m. 3*, Bloomington #25174); Act5C-Cas9 (Port et al., 2014) (Bloomington # 54590); *ct*-Gal4(Pfeiffer et al., 2008) (Bloomington # 45603); *hth*^NP5332^-Gal4 (DGRC Kyoto #104957); UAS-RNAi^ct^ (VDRC #v5687); UAS-RNAi^ct^ (VDRC #v4138); UAS-RNAi^luc^ (Bloomington #31603); *D.m.ey3.5*^*G*^*GFP* (this study); *D.m.ey3.5*^*A*^*GFP* (this study); *D.m.ey3.5*^*G*^*mCherry* (this study); *D.m.ey3.5*^*A*^*mCherry* (this study); *D.m.ey3.5*^*NoCt*^*GFP* (this study); *D.m.ey3.5*^*ConsensusCt*^*GFP* (this study); CRISPR^WT^ (this study); CRISPR1 (this study); CRISPR2 (this study); CRISPR3 (this study); CRISPR4 (this study). *D. ananassae*: isofemale stocks from Kisangani, Congo (DSSC 14024-0371.30) and Mumbai, India (DSSC 14024-0371.31); *D. yakuba*: stocks from Ivory Coast (DSSC 14021-8209.0261.00) and Liberia (Reference Genome strain DSSC 14021-0261.01). *D. pseudoobscura*: isofemale stocks from Catalina Island, California, USA (Cat; DSSC 14011-0121.121) and Chiracahua Mountains, Arizona, USA (DSSC 14011-0121.118). *D. virilis*: isofemale stock from Gikongoro, Rwanda (DSSC 15010-1051.118). All stocks were maintained on standard cornmeal diet food, at 25°C except where mentioned otherwise.

### Developmental stages

Correspondence of developmental stages between *D. mel*. and *D. pse*. was determined based on developmental transitions – larval molts, pupa formation - and morphological features – morphology of the embryo, numbers of rows of differentiated photoreceptors. Staging was performed at 25°C. Embryos were collected on grape fruit plates complemented with yeast paste exchanged every 2 hours. Freshly hatched L1 larvae were collected every two hours and transferred to corn meal food vials in a density control fashion (20-25 larvae / vial).

### *Drosophila melanogaster* Genome Assembly

All *D. mel*. genome positions refer to BDGP release 6 assembly (GCA_000001215.4).

### Constructs

Enhancer reporter constructs were generated using the Gateway Recombination Cloning Technology (ThermoFischer Scientific). *D. pse. ey3.3* and *D. mel. ey3.5* regulatory sequences were amplified respectively from *D. pse*. (from stock Cat; DSSC 14011-0121.121), Hikone-AS (for the *G*-variant) and Canton-S^BH^ (for the *A*-variant) genomic DNA (extracted using Qiagen DNeasy Blood and Tissue Kit #69504) and cloned into the Gateway pDONR221 entry vector (ThermoFischer Scientific #12536017) following the provider specifications. Primers for the enhancer amplifications are:

#### pEntry-ey^33Pse^

*forward*:GGGGACAAGTTTGT ACAAAAAAGCAGGCTAAGT GGT AGT GGACT AGG and

*reverse*:GGGGACCACTTTGTACAAGAAAGCTGGGTCCTAGAATTTTGCTAACGC;

#### pEntry-ey^35CSBH^ and pEntry-ey^35Hik^

*forward*:GGGGACAAGTTTGTACAAAAAAGCAGGCTGGACTAGGCGGTATTGCT and

*reverse*:GGGGACCACTTTGTACAAGAAAGCTGGGTTTTGCTCACACATCCATTTG.

The entry vectors with mutated forms of *ey3.5* enhancer, pEntry-ey^3.5NoCt^ and pEntry-ey^3.5consensusCt^ were generated by modifying the pEntry-ey^3.5Hik^ using primers carrying the corresponding mutations. These primers were (mutated nucleotides are in capital letters):

#### pEntry-ey^3.5NoCt^

*forward*: caataaaatggttgg**C**a**G**tttttcgaactttcg

*reverse*: cgaaagttcgaaaaa**C**t**G**ccaaccattttattg

#### pEntry-ey^3.5consensusCt^

*forward*: taaaatggtt**T**gaactttttcgaactttcg

*reverse*: gaaaaagttc**A**aaccattttattgttttc

Enhancer inserts were next transferred using Gateway recombination into mCherry- and GFP-expressing enhancer reporter vectors amenable to phiC31 integration ‒mediated transgenesis (Aerts et al., 2010; Quan et al., 2016).

pU6gRNA^*ey*^: the following complementary phospho-oligomers were used to generate a double strand DNA sequence encoding the *ey* eye-enhancer guide RNA (gRNA): *forward*: phospho-CTTCGTCGAAAACAAT AAAAT GGT; *reverse*: phospho-AAACACCATTTTATTGTTTTCGAC. After hybridization, the resulting double-strand DNA was cloned into the pU6-BBS1-chiRNA plasmid (Addgene #45946).

### Transgenic lines

Transformant flies carrying enhancer reporter constructs were generated by BestGeneInc. All constructs were integrated at the Attp2 landing site using phiC31 recombination.

### Scanning electron microscopy

Whole flies were fixed overnight at 4°C in a 1:1 mix of 4% formaldehyde in phosphate buffer pH 7.2 and 100 % ethanol and dehydrated successively in graded ethanol series, hexamethyldisilazane (HMDS) and in a dessicator. Fly heads were mounted on specimen studs using silver paint in two distinct orientations: dorsal head up (for whole head imaging) and lateral (for ommatidia counts and measures). Samples were subsequently coated with platinium and images acquired in LEI mode with a JEOL JSM 7401F microscope at magnifications ranging from 120 (heads overviews) to 1900 times (ommatidia width) (Schneider et al., 2012).

### Transmitted light and confocal microscopy

Preparations of adult heads for the acquisition of light microscopy images were acquired from non-fixed, freshly cut adult heads glued laterally on glass slides. Images were acquired using a camera DFC295 (Leica) mounted on a DMRXA (Leica) microscope, operated via the open-source software Micro-Manager (Edelstein et al., 2014). Fluorescent preparations of embryos and imaginal discs were acquired using a Nikon A1R Eclipse Ti, a Leica TCS SP5 II or a Leica SP8 confocal microscope operated by the accompanying company software.

### Behavioral measure of visual acuity

The experimental set-up is modified from (Buchner, 1976). It exploits the spontaneous tendency of fruit flies to adjust their trajectory to the surrounding landscape. Presented with rotating vertical black stripes, tethered flies spontaneously follow their movement. Narrowing the angular distance between the stripes beyond its resolving capacities makes the fly move in the opposite direction, due to an interference phenomenon similar to what we perceive when looking at the wheels of a starting train. It consists of two tracking balls and of two computer screens on which moving vertical black and white stripes are displayed in a window of 90° horizontal and 74° vertical extension. The width of the stripes (spatial wavelength λ) as they move on the flat screen are adjusted such that they subtend a constant angle as seen from the fly 40 mm away of the screen. Pattern speed w is adapted to maintain the “contrast frequency” at 1 Hz. A positive optomotor response indicates the tendency of the flies to follow the direction of the movement of the stripes. Reduction of λ below the resolving power of the eye causes an inversion of the apparent direction of the movement of the stripes due to the geometrical interference between the fly’s vertical columns of ommatidia and the vertical stripes, and is accompanied by the inversion of the fly response towards negative values. The λ value at which the response of the fly is inverted (zero-crossing angle, 2ΔΦ) provides a measure of spatial resolution (or visual acuity). Recordings of female *D. mel*. (Canton-S; n=9) and *D. pse*. (Cat; n=9) were performed simultaneously, with alternating assignments. We calculated the zero-crossing and its variance from the two average responses surrounding the zero-crossing (one positive, one negative) using linear interpolation and error propagation followed by t-test for differences between 2 means.

### Eye and head measurements

All head and eye measurements were performed on female flies using ImageJ (Schneider et al., 2012). Adult Eye/Head ratio was expressed as 100 × (Head width (HW) − Face width (FW)) / HW (Figure 1). Ommatidia width was measured on high magnification SEM images as the distance between one interommatidial bristle and the opposite angle of the facet. For each sample, measures of six adjacent ommatidia localized at the center of the eye were taken. To limit underestimation of the ommatidia width due to perspective projection distortion, samples were carefully oriented prior to image acquisition.

### Ommatidia numbers

Ommatidia numbers were manually counted on SEM images using the Image J plugin “Cell counter”. We also developed an alternative method based on the approximation of the compound eye to an ellipse. With this method, the ommatidia number is calculated as the surface of an ellipse whose large and small axes correspond to the numbers of ommatidia along the compound eye anterior-posterior and dorso-ventral axes (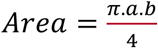 with a and b as the lengths of the large and small ellipse axes; Fig. S4). This method accommodates lower resolution images and does not require the use of SEM. Bland-Altman method (Bland and Altman, 1986) was used to compare the outcome of the two methods applied on a common set of SEM images. The ellipse method results in an overestimation of approximately 20 ommatidia as compared to the manual counting (bias mean = 20. 29; SD = 11. 70). Importantly, this difference is independent of ommatidia number (Figure S4). Facet number estimation of Cut RNAi and CRISPR flies (Fig. 3b and d) were performed blind regarding to the genotype.

### Immunostainings

Fixation and immunostainings were performed following standard procedures as described in (Patel, 1994) (embryos) and in (Blair, 2000) (imaginal discs). We used the following primary antibodies: mouse anti-Ct (1:10, DHSB hybridoma supernatant 2B10, deposited by Rubin G. M.), mouse anti-Eya (1:75, DHSB hybridoma supernatant eya10H6, deposited by Benzer, S. and Bonini, N.M.), rat anti-elav (1:100, DHSB hybridoma supernatant Rat-Elav-7E8A10, deposited by Rubin G. M.), rat anti-Ey (1:300, received from P. Callaerts); anti-phosphorylated Histone 3 (1:1000, pSer10; Merck Millipore #382159); sheep anti-atonal (1:1000, received from A. Jarman and P. zur Lage); mouse anti-GFP 3E6 (1:1000, Invitrogen, #A11120); rabbit anti-DsRed (1:1000, Clonetech, #632496). Secondary antibodies conjugated with Alexa 488, Alexa 555 and Alexa 647 were used at 1:200 concentration (Molecular Probes). Nuclei were counterstained using Draq-5 (1:1000 in PBS, Abcam). All samples were mounted in Vectashield (Molecular Probes).

### Measurements on embryonic and larval eye-antennal discs

Numbers of Ey-positive embryonic eye-antennal disc cells were counted manually. To measure the surface of the larval eye-antennal disc and eye progenitor field, regions of interest were selected manually using the Image J freehand selection tool. The number of mitotic pH3-positive cells was automatically counted using the Dead-Easy Mito-Glia ImageJ Plugin (Forero et al., 2010). The mitotic index was calculated as the number of mitotic cells per surface of the Eya-positive eye progenitor field.

### Quantification of mCherry and GFP colocalization

Protocol for pixel-based quantifications of mCherry and GFP colocalization was adapted from (Oliva et al., 2016). We used Fiji/ImageJ2/ImgLib2 (Pietzsch et al., 2012; Rueden et al., 2017; Schindelin et al., 2012) macro implemented in Jython. Raw images were imported using BioFormats library (Linkert et al., 2010). EADs were manually segmented in each stack by the user. Stack threshold levels for each channel were calculated using preselected autothresholding algorithms available in Fiji (Huang for both channels). Determined threshold levels were used to calculate Mander’s overlap coefficient using Fiji implementation of the colocalization algorithm. Code for the macro is available on GitHub (https://github.com/rejsmont/FijiScripts/blob/master/mColoc3D.py). The proportion of pixels expressing solely mCherry is used as a measure for GFP retraction.

### DNA sequence alignments

Pairwise alignement of the D.pse and D.mel. intronic eye enhancers was performed with BLAST (McGinnis and Madden, 2004)(blast.ncbi.nlm.nih.gov) as described in (Swanson et al., 2011). The 22 *%* of alignment corresponds to “query coverage “in blastn output and indicates the amount of D. mel. intron sequence included in blocks aligned to the *D. pse* ortholog by BLAST.

### Population genetics

We compiled whole genome sequencing data from multiple geographic samples collected in Africa, Europe, North America, Asia and Australia to investigate worldwide allele frequency patterns of the *eyeless* SNP at position Chr 4: 710326 in natural populations. This dataset consisted of single individual sequencing data (Campo et al., 2013; Grenier et al., 2015; Lack et al., 2015; Lack et al., 2016; Langley et al., 2012; Pool et al., 2012) and Pool-Seq data from various sources (Bastide et al., 2013; Bergland et al., 2014; Kapun et al., 2016; Orozco-terWengel et al., 2012; Reinhardt et al., 2014) (Table S2). For single individuals, we obtained genotypes of the focal SNP from the *Drosophila* Genome Nexus website (DGN; http://www.johnpool.net/genomes.html) and estimated allele frequencies based on the number of chromosomes carrying the *A*-variant for populations with at least ten sequenced individuals. For Pool-Seq datasets, we re-mapped quality-filtered raw data as described in (Kapun et al., 2016) and estimated allele frequencies based on read counts of the *A*-variant relative to the total coverage. To increase sequence coverage in Queensland and Tasmania, we merged libraries of multiple collections at the corresponding locations (Reinhardt et al., 2014). We further used a collection of Pool-Seq data from 48 population samples collected across Europe by the DrosEU consortium (Kapun et al., 2018) (accession number: PRJNA388788) for an in-depth analysis of spatial distribution of the *A*-variant. Specifically, we tested for clinal distribution along the latitudinal and longitudinal axes by means of generalized linear models (GLMs) in *R* based on allele counts to account for the biallelic nature of the focal SNP. We further contrasted the clinality of 4: 710,326 to 21,008 putatively neutral genome-wide SNPs located in short introns (<60bp) and in distance to chromosomal inversions (Clemente and Vogl, 2012; Parsch et al., 2010). To test if the observed *p*-value from a GLM at the focal SNP deviates from neutral expectation we empirically assessed significance. We therefore generated empirical cumulative density functions (ECDF) based on the −log10 transformed *p*-values of all neutral SNPs and calculated the area under the ECDF confined by the −log10 *p*-value of the focal SNP and the upper tail of the distribution by integration. This area corresponds to the percentile of neutral SNPs with *p*-values equal or smaller than the focal SNP and thus summarizes the significance of clinality for 4:710,326 relative to genome-wide neutral estimates. We further characterized chromosome-wide patterns of genetic variation by estimating the population genetics statistics *π* and Tajima’s *D* for all 48 samples from the DrosEU dataset using Pool-Gen (Kapun et al., 2018) with implemented corrections for Pool-Seq data (Futschik and Schlotterer, 2010; Kofler et al., 2011). At last, we tested whether very rare occurrences of the *A*-variant in Sub-Saharan Africa may be due to admixture with non-African genetic variation. We therefore used admixture probability estimates from (Lack et al., 2015) (see Table S2) to classify African lines as admixed (>10% of the autosomes admixed) or non-admixed (≤ 10% of the autosomes admixed) and compared genotype counts for admixed and non-admixed lines by means of Fisher exact tests (FET) in *R*.

### CRISPR/Cas9 SNP engineering

For editing the *ey* eye enhancer, we injected SNP^G^ homozygous *D. mel*. Act5-Cas9 embryos(Port et al., 2014) with two constructs respectively encoding the guide RNA (pU6gRNA^*ey*^) and the SNP^A^-carrying *ey* eye enhancer sequence (pEntry-ey^3.5CSBH^), each of them at a concentration of 500 ng/μl (Port et al., 2015). Candidates were screened using allele-specific PCR. We isolated one CRISPR modified male from which we established four CRISPR SNP^A^ lines. In parallel, a control line was established by mating non-injected Act-Cas9 flies following the same scheme their injected siblings. Sequencing the *ey* eye enhancer from the transformed SNP^A^ and of the non-injected SNP^G^ control stocks confirmed that they were differing only by this single nucleotide.

### Allele-specific PCR for screening CRISPR alleles

SNP^A^ and SNP^G^ alleles were detected by allele-specific PCR using a common reverse primer (Ey-R3: AGAAATATCACATGGCCGAG) and one of two specific forward primers differing by the 3’ most nucleotide (either A or G) and including a mismatch (underlined) to increase binding specificity (Ey-SNP^G^-F: GGAATCGAAAACAATAAAATGG**<u>C</u>**TG**G**; Ey-SNP^A^-F: GGAATCGAAAACAAT AAAATGG**<u>C</u>**TG**A**).

### Image Processing

Except mentioned otherwise, all image processing was performed using Image J (versions 1.45 to 1.48)(Schneider et al., 2012).

### Sample sizes

A fixed sample size was chosen in all cases prior to phenotypic analyses, samples were excluded exclusively prior to quantification based on image quality or developmental stage, leading to differences between groups. Except where stated otherwise, all statistical tests and charts were performed using Graphpad Prism 7 (Graphpad software Inc.). Details on the tests, sample sizes and p values are indicated in figure legends except where mentioned otherwise.

## ACKNOWLEDGMENTS

This work was funded by VIB (B.A.H), the Belspo WiBrain Interuniversity Attraction Pole network and Fonds Wetenschappelijke Onderzoeks (FWO) grant G.0503.12 (B.A.H.). Work in B.A.H lab is currently funded by the Institut Hospitalier Universitaire (IHU) and the Institut du Cerveau et de la Moëlle Epinière (ICM), Paris, France. S.W. was a fellow of the FLiACT Marie Curie ITN (FP7, EU). M.K. and T.F. were supported by a Swiss National Science Foundation (SNSF) grant (PP00P3_133641) to T.F. We thank D. Pottier, S. Villain, R. Ejsmont, N. Mora, A. Soldano, N. Gompel, N. Posnien and D. Nunes for advices, discussion or/and technical help; J. Clements and P. Callaerts for sharing *Drosophila* strains and antibodies. We acknowledge the *Drosophila* Species Stock Center (DSSC), the Bloomington Stock Center (NIH P40OD018537), the Kyoto Stock Center (DGRC), for sending *Drosophila* stocks, the Developmental Studies Hybridoma Bank (DHSN, University of Iowa) for sending antibodies and Addgene for sending plasmids. We are grateful to P. Baatsen from the Electronic Microscopy Platform for support with SEM sample preparation and imaging. Confocal imaging was performed at the VIB Light Microscopy and Imaging Network (Limone) and at the Plateforme d’imagerie cellulaire (PICPS) at the ICM. The Leica TCS Sp5 II confocal microscope and the Nikon were acquired respectively by the VIB BioImaging Core and through a Hercules type 1 grant (AKUL/09/037) to W. Annaert. B.A.H. is an Allen Distinguished Investigator and an Einstein Visiting Fellow of the Berlin Institute of Health.

## AUTHOR CONTRIBUTIONS

A.R. conceived the project. A.R. and B.A.H. designed the experiments. A.R. and J.Y. acquired SEM and light microscopy images. A.R. and S.W. performed and imaged immunostainings on imaginal discs, A.R. quantified and analyzed the imaging data. A. R. and A.C. generated the constructs and performed allele-specific PCR. J.Y. injected the CRISPR constructs. E.B. and R.W. performed and analyzed the optomotor tests. M. K. and T. F. performed population genetics analyses. A.R. and B.A.H. wrote the manuscript.

## Supplemental Information

**Figure S1.**
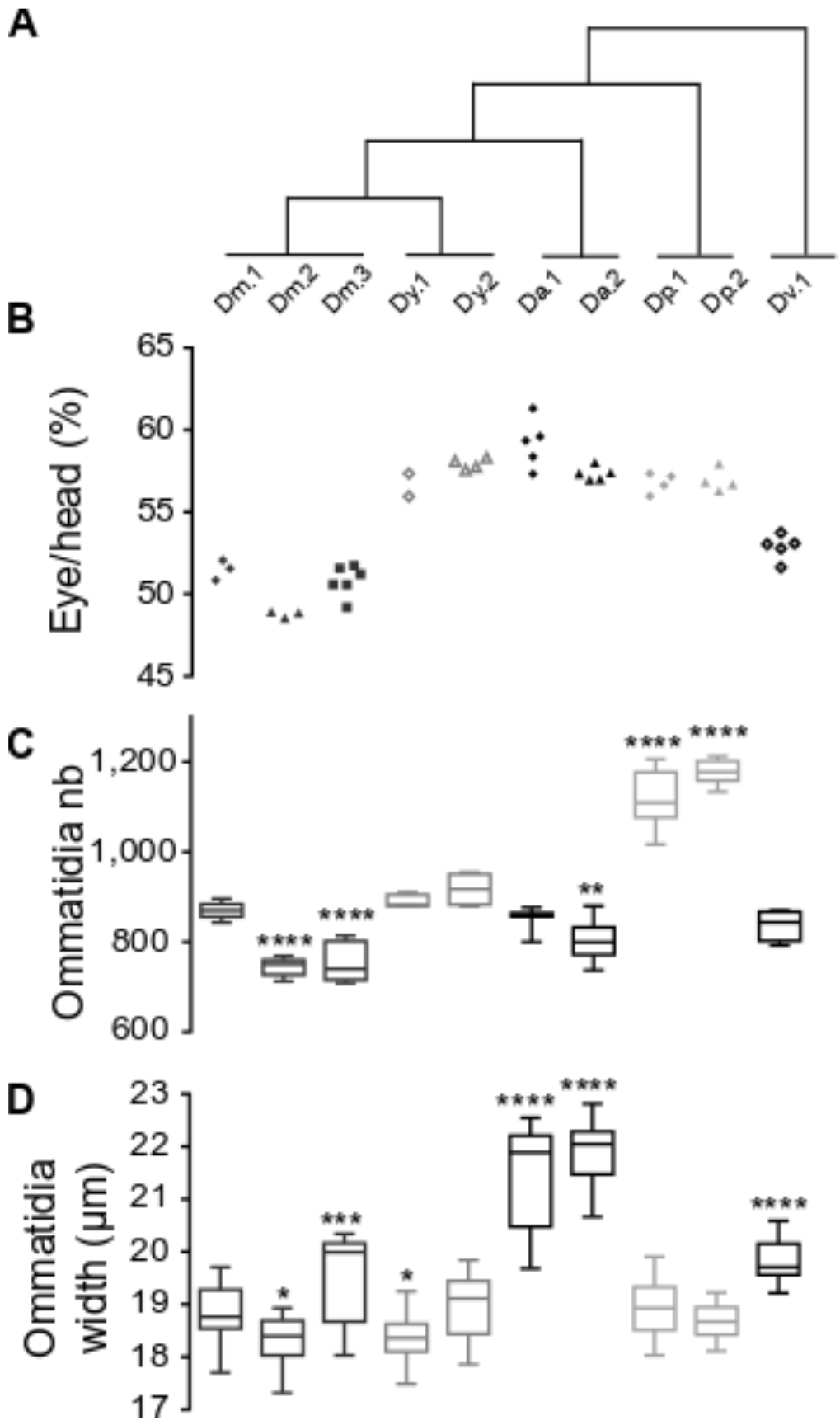
Natural variation in eye size in *Drosophila*, Related to Figure 1. (A to C’) Eye size comparison between five *Drosophila* species: *D. melanogaster* (*D. mel*.), *D. yakuba* (*D. yak*.), *D. ananassae* (*D.ana*.), *D. pseudoobscura* (*D. pse*.), *D. virilis* (*D. vir*.). Different numbers indicate different strains (*see Methods*). Boxes indicate interquartile ranges, lines medians and whiskers data ranges. (A) Phylogenic relationship between the five species (tree branches are not scaled). (B) Eye/head width ratio. (C) Ommatidia number counted on SEM images. Ordinary one-way ANOVA **** *p*<*0.0001* followed by Dunnett’s multiple comparisons. See also Table S1. (D) Ommatidia width. Ordinary one-way ANOVA **** *p*<*0.0001* followed by Dunnett’s multiple comparisons. See also Table S1

**Figure S2.**
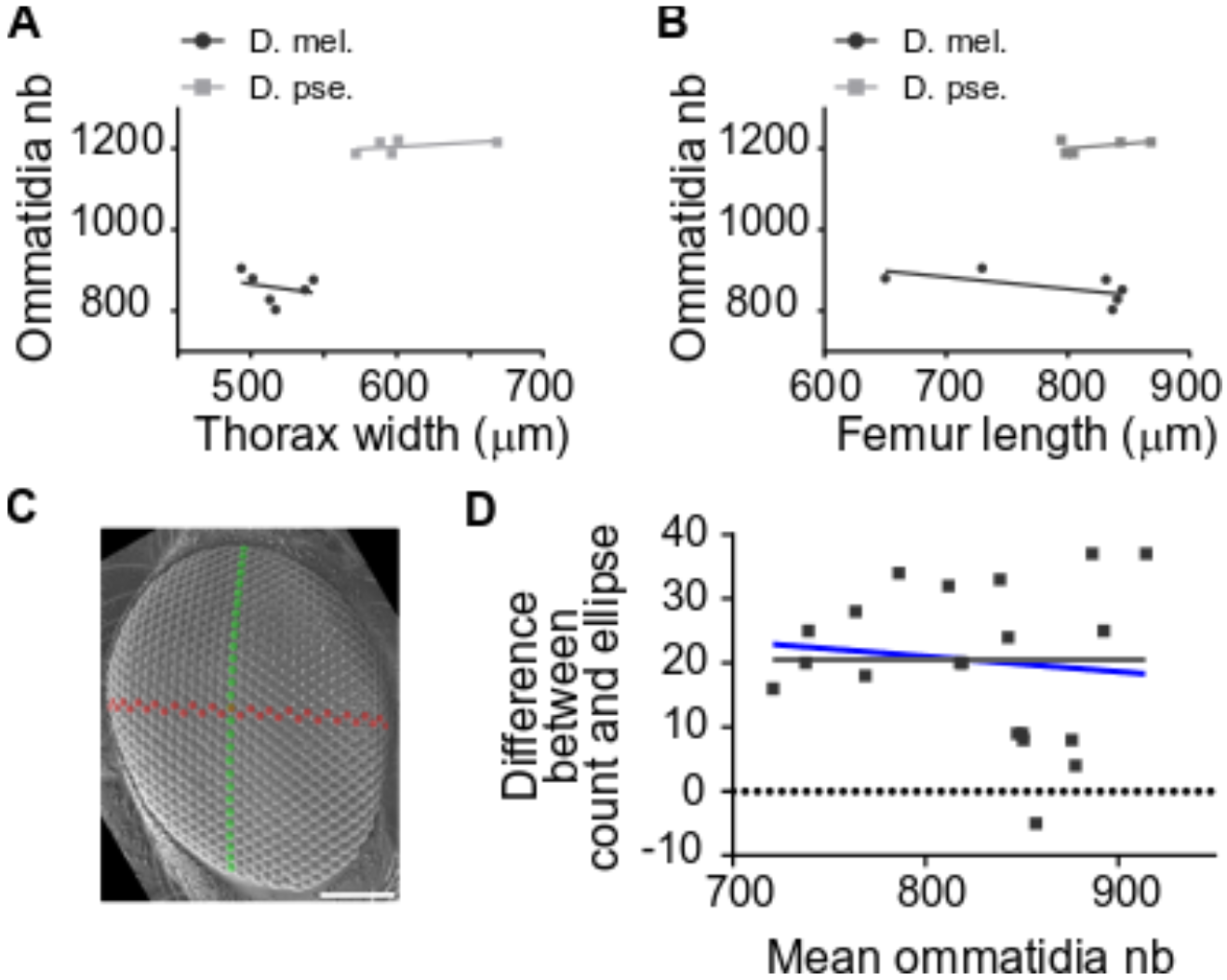
Ommatidia number variation: scaling and methods, Related to Figures 1, 4 and 6. (A) *D. mel*. and *D. pse*. ommatidia numbers in relation to body size estimated from thorax width. Intercepts (F=84.62, Dfn=1, Dfd=8, p<0.0001) but not slopes (F=0.695, Dfn=1, Dfd=7, p=0.4320) are significantly different indicating that interspecific differences in ommatidia numbers cannot be solely explained by differences in body size, as estimated by thorax width. (B) *D. mel*. and *D. pse*. ommatidia numbers in relation to body size estimated from mesothoracic femur length. Intercepts (F=419.2, Dfn=1, Dfd=8, p<0.0001) but not slopes (F=1.454, Dfn=1, Dfd=7, p=0.2671) are significantly different indicating that interspecific differences in ommatidia numbers cannot be solely explained by differences in body size, as estimated by femur length. (C) SEM image of a *D. mel*. Hikone-AS eye. *Green*: dorso-ventral axis; *Red*: anterior-posterior axis. Scale bar: 100 μm. (D) Bland-Altman chart plotting the difference in ommatidia number measured by two methods (ellipse-based estimation vs direct counting) over their mean (Bland and Altman, 1986). Comparison of fits indicates that the difference between the two measurements is independent of the mean (null hypothesis, grey line: slope= 0.0; alternative hypothesis blue line: slope unconstrained = −0.02372; *p*=*0.6212*).

**Figure S3.**
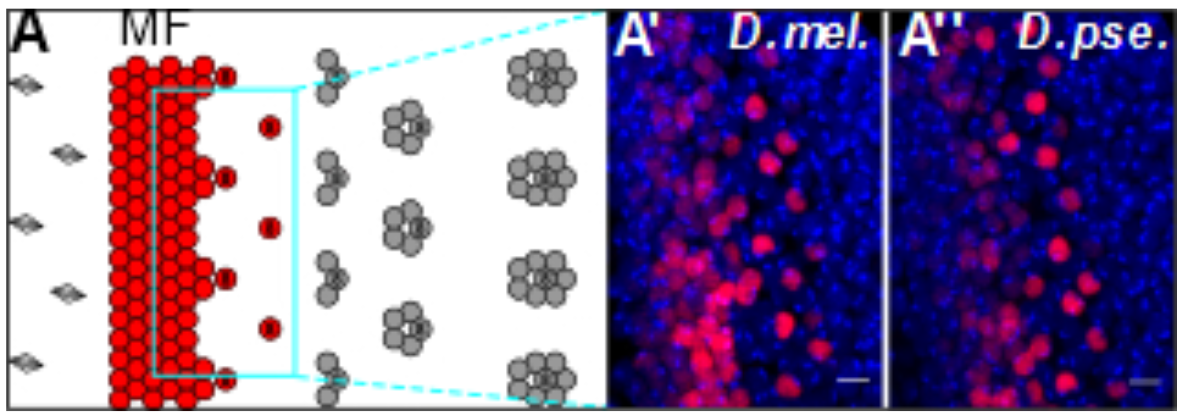
Developmental origin of eye size variation in *D. mel*. and *D. pse*., Related to Figure 2. (A) Schematics of the first steps of retinal differentiation showing the singling-out of committed Ato-expressing R8 ommatidia progenitor cells and subsequent steps of ommatidia assembly. (A’ and A”) The density of Ato-expressing R8 progenitors (in red in A’ and A”) is similar in the two species. *Red*: anti-Ato immunostaining; *blue*: DAPI. Anterior is at the left. Scale bars: 5 μm.

**Figure S4.**
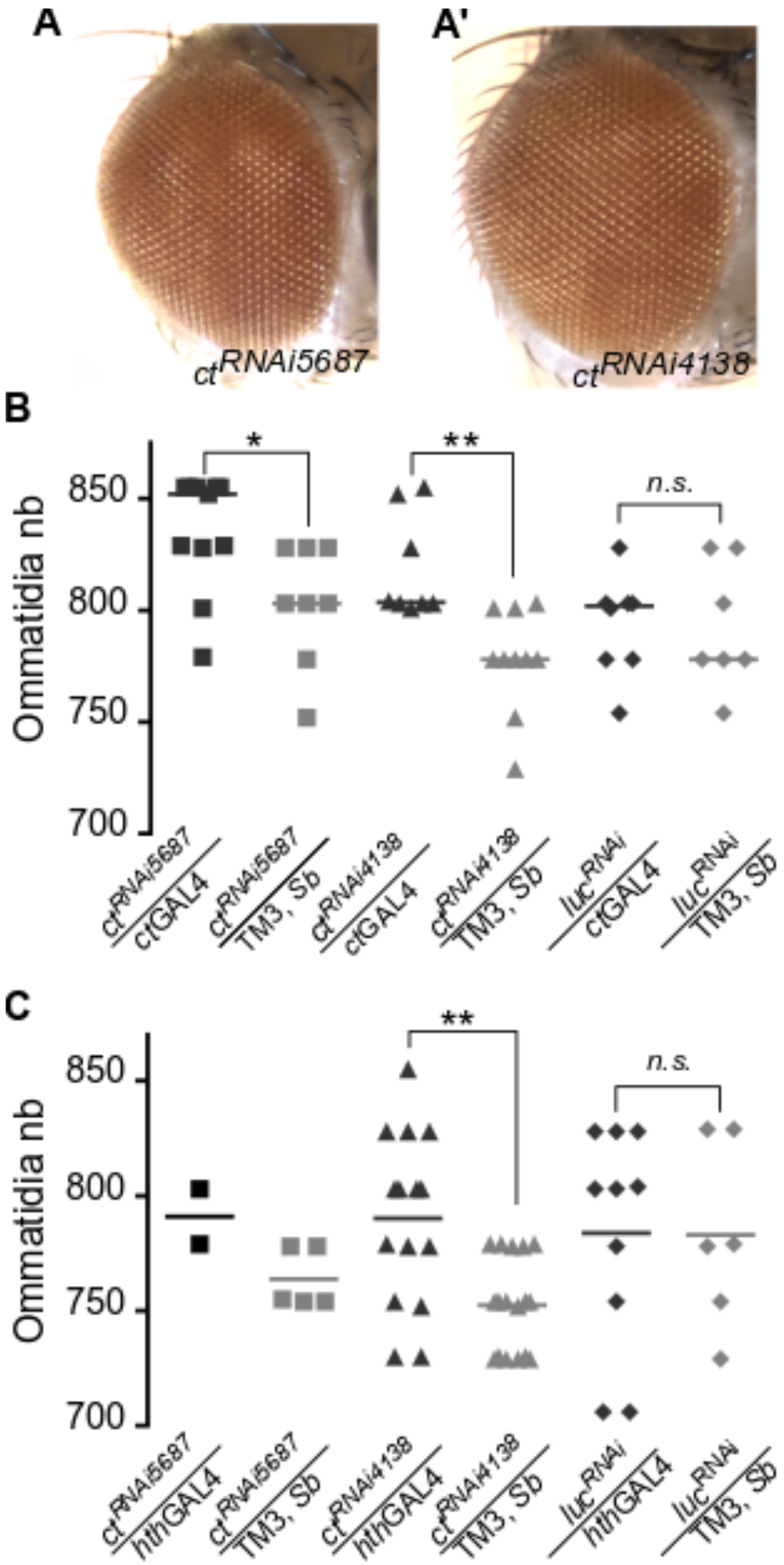
RNAi-mediated KD of *ct* increases facet number, Related to Figure 4. (A and A’) RNAi-mediated KD of *ct* using two distinct RNAi constructs does not induce gross morphology defects in the compound eye. Gal4 driver: *ctGal4*. (B) Overexpression of two UAS-*ct*^*RNAi*^ and one UAS-*luciferase*^*RNAi*^ constructs under the control of *ct*GAL4.Sample sizes from left to right (n=23, n=13, n=8, n=10, n=8, n=7, n=6). Ordinary one-way ANOVA **** *p*<*0.0001* followed by Sidak’s multiple comparisons: *ct*^*RNAi5*687^/*ctGal4* vs *c*^*tRNAi5*687^/ TM3, Sb \*\*\*\**p*<*0.0001*; *ct*^*RNAi5*687^/*ctGal4* vs *ctGAL4*/ + **** *p*<*0.0001*; *ct*^*RNAi4138*^/*ctGal4 vs ct*^*RNAi4138*^/TM3, Sb \**p*=*0.0126*; *ct*^*RNAi4138*^/*ctGal4 vs ctGAL4*/ +**** *p*<*0.0001*; *luc*^*RNAi*^/*ctGal4* vs *luc*^*RNAi*^ / TM3, Sb *n. s. p*>*0.9999*; *luc*^*RNAi*^ /*ctGal4 vs ctGAL4*/ + ** *p*=*0.0036*. (C) Overexpression of two UAS-*ct*RNAi and one UAS-*lucferase*RNAi constructs under the control of *hth*GAL4. Sample sizes, from left to right (n=2, n=5, n=15, n=17, n=10, n=7). Sample size for ctRNAi5687/hthGAL4 was low due to the lethality or gross morphological defects caused by this allelic combination. Ordinary one-way ANOVA ** *p*=*0.0089* followed by Sidak’s multiple comparisons: *ct*^*RNAi4138*^/*hthGal4 vs ct*^*RNAi4138*^/TM3, Sb ** *p*=*0.0047*; *luc*^*RNAi*^ /*hthGal4* vs *luc*^*RNAi*^ / TM3, Sb *n. s. p*=0*.2152*. (B and C) Scatter dot plots. Line indicates the mean.

**Figure S5.**
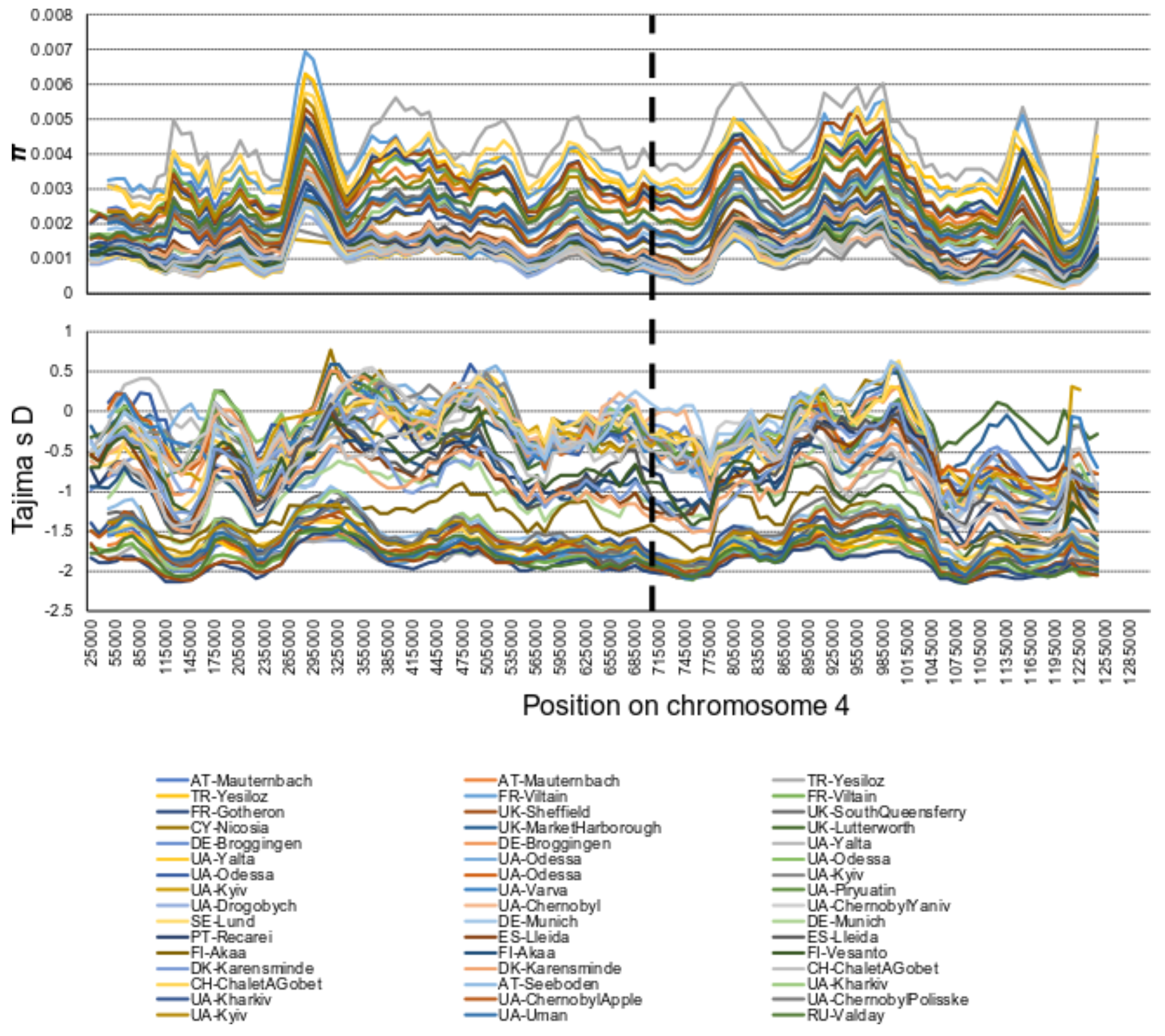
Genetic variation of the fourth chromosome in Europe, Related to Figure 5. The distribution of *π* (top panel) and Tajima’s *D* (bottom panel) in 50kb windows with 10kb step-size for 48 population samples from Europe. The vertical dashed black line indicates the approximate genomic position of the focal SNP at position Chr 4: 710326.

**Table S1.** Natural variation in *Drosophila* eye size, Related to Figure S1.

| Genotype | Ommatidia Number |  | Ommatidia width |  |
| --- | --- | --- | --- | --- |
|  | **n** | ***adjusted p value*** | **n** | ***adjusted p value*** |
| *D. m. 2* | n=7 | *p = 0.0001* | n=24 | *p = 0.0161* |
| *D. m. 3* | n=8 | *p = 0.0001* | n=24 | *p = 0.0002* |
| *D. y. 1* | n=5 | *p = 0.9396* | n=24 | *p = 0.0462* |
| *D. y. 2* | n=4 | *p = 0.2764* | n=24 | *p = 0.9922* |
| *D. a. 1* | n=8 | *p = 0.9770* | n=24 | *p = 0.0001* |
| *D. a. 2* | n=8 | *p = 0.0083* | n=24 | *p = 0.0001* |
| *D. p. 1* | n=9 | *p = 0.0001* | n=24 | *p = 0.9994* |
| *D. p. 2* | n=8 | *p = 0.0001* | n=24 | *p = 0.9072* |
| *D. v.* | n=4 | *p = 0.6782* | n=24 | *p = 0.0001* |

Sample sizes and results of Dunnett’s multiple comparison tests following ordinary one-way ANOVA from Figure S1. Comparisons towards *D. m. 1* (Canton-S^BH^; ommatidia number sample size n=6; ommatidia width sample sizes n=24).

**Table S2. Worldwide allele frequency patterns, Related to Figure 5**

*Data are presented in a separate Excel document*.

Origin, data type, data source and allele frequencies of the *A*-variant of the focal SNP at position Chr 4: 710326 of world-wide populations with sample sizes ≥ 10 individuals.

**Supplementary Table 3. Isofemale line genotypes, Related to Figure 5**

*Data are presented in a separate Excel document*.

Genotypes and admixture status for isofemale lines from Sub-Saharan Africa.

